# Gram-positive bacteria have different susceptibilities to the host-derived antimicrobial lysophosphatidylcholine and adapt to lysophosphatidylcholine-induced stress via distinct mechanisms

**DOI:** 10.64898/2026.11.17.767358

**Authors:** Yahan Wei, Ziqiang Guan, Chahat Upreti, Savannah E. Pas, Julie Nguyen, Priya M. Christensen, Gayatri Chandrasenan, Lily E. Tillett, Guan H. Chen, Kelli L. Palmer

**Affiliations:** School of Podiatric Medicine, The University of Texas Rio Grande Valley, Harlingen, TX, USA; Department of Biochemistry, Duke University School of Medicine, Durham, North Carolina, USA; Broad Institute of MIT and Harvard, Cambridge, Boston, MA, USA; Department of Biological Sciences, The University of Texas at Dallas, Richardson, TX, USA

**Keywords:** lysophosphatidylcholine, plasmalogen, Gram-positive bacteria, antimicrobial lipids

## Abstract

The human lipid metabolites, lysophosphatidylcholine (LPC) and plasmenyl-LPC (pLPC), modulate both the metabolic health status and innate immunity of an individual. Previous studies reported direct antibacterial activity of LPC, while pLPC has not been evaluated. Meanwhile, some bacteria harvest LPC for producing phosphatidylcholine (PC); others produce plasmalogens under anaerobic conditions. These discoveries suggest that both LPC and pLPC can mediate host-microbe interactions. In this study, we demonstrated diverse susceptibility profiles of human pathobionts to LPC(18:0) and pLPC(18:0) at physiologically-relevant concentrations. Though the Gram-negative bacteria *Escherichia coli* and *Pseudomonas aeruginosa* were not susceptible to LPC or pLPC, all tested Gram-positive bacteria, including *Staphylococcus aureus*, *Streptococcus agalactiae*, *Enterococcus faecalis*, and *E. faecium* were susceptible to pLPC, suggesting robust antimicrobial activity of this lipid *in vivo*. While LPC can effectively kill *S. aureus* and *S. agalactiae*, it does not completely inhibit the growth of enterococci. Moreover, pretreatment of LPC-containing medium with *E. faecalis* rescues *S. agalactiae* from LPC killing, suggesting that *E. faecalis* deactivates LPC extracellularly and can protect co-resident LPC-sensitive pathobionts. Finally, we observed the emergence of spontaneous LPC-resistant *S. aureus.* These strains have point mutations in sensor kinase gene *walK* and the lipid metabolism genes *fabF* and *farR*, which lead to increased expression of the efflux pump gene *farE*. Collectively, these results support the antibacterial effects of LPC and pLPC and indicate distinct bacterial adaptation mechanisms, providing novel insights on lipid-mediated host-bacterial interactions.

**Importance:** Bacteria encounter host lipid metabolites during infections. Prior work established that the human lipid metabolite LPC has antibacterial activity, yet some bacteria harvest LPC for producing PC. In this study, we evaluated LPC susceptibility in several major human pathobiont species and investigated their mechanisms for LPC stress response. We additionally investigated the LPC variant pLPC, which has not been previously investigated for antimicrobial activity. We found different LPC-resistance mechanisms in the Gram-positive bacteria *E. faecalis* and *S. aureus*, neither of which produce PC from LPC. We additionally identified that pLPC has robust antibacterial activity against Gram-positive bacteria. All together, these results demonstrate the direct antibacterial activities of human lipid metabolites LPC and pLPC. LPC and pLPC are biomarkers whose plasma levels change in response to an individual’s health status. Our findings suggest a role for these lipid metabolites in shaping the composition and genetic diversity of the human microbiota.

## Introduction

Human lipid metabolites modulate both the metabolic health status and innate immunity of an individual, both of which affect infection risks.^1,2^ Lipid metabolites have different distributions among organs and tissues, which is strongly associated with their primary biological functions.^2–4^

One such example is the lipid species that contributes to about 20% of total phospholipids in humans, the plasmalogens.^5^ A plasmalogen contains a vinyl ester bond that links an acyl chain with its glycerol backbone. The most common polar headgroups of plasmalogens at the sn-3 position of the glycerol are ethanolamine and choline.^5^ Plasmalogens are primarily enriched in the brain (constitutes 76 to 100% of phospholipids in white matter^5^) and heart (up to 42% of total phosphatidylcholines^6^). As such, they are strongly linked with both cardiovascular and neurological health,^3^ with decreased plasma levels identified as biomarkers for both cardiovascular and neurodegenerative diseases.^3,7^ Additionally, due to the existence of an unsaturated double bond, plasmalogens have protective roles against oxidation, and thus modulate oxidation-associated inflammation.^7^

Another example is lysophosphatidylcholine (LPC), a lipid metabolite that is prevalent in tissue and in circulation.^8^ LPC is mainly produced by the hydrolysis of phosphatidylcholine (PC), the primary lipid component of eukaryotic cell membranes, and thus signals tissue damage and induces immune responses.^8^ On the other hand, as a component of modified low-density lipoprotein (LDL) and oxidized LDL, LPC is also strongly linked with cardiovascular and neurodegenerative diseases, with fluctuating plasma levels under different health conditions.^2,8^ Particularly, LPC can induce glucose uptake by adipocytes and is observed with a decreased serum level in people with Type 2 Diabetes,^9–11^ a metabolic disorder that is associated with increased infection risks and significantly more severe infection consequences.^12,13^

In addition to modulating immune responses, human-derived lipid metabolites have direct antimicrobial properties. LPC sensitizes both Gram-negative and Gram-positive bacteria to antibiotics,^14–16^ lyses cell-wall-deficient bacteria including mycoplasmas and L-form *Staphylococcus aureus*, and induces autolysis in *Bacillus subtilis*.^17^ Additionally, administration of LPC in combination with antibiotics can cure bacterial sepsis in a murine infection model.^16,18,19^ Yet, the underlying antimicrobial mechanism of action of LPC is undefined. It is suggested that LPC kills mycobacteria via the free fatty acids (FFAs) released through its hydrolysis.^20^ FFAs, particularly those with medium- (8 to 12 carbons) and long-chain (more than 12 carbons) lengths and containing one or more unsaturated double bonds have potent antibacterial activities against Gram-positive bacteria.^21–24^ The primarily proposed antibacterial mechanism of FFAs is the disruption of membrane integrity, which results in cell lysis, as indicated by the leakage of low-molecular-weight proteins from *S. aureus* caused by palmitoleic acid (C16:1Δ9);^22^ and the lysis of *Enterococcus faecalis* induced by oleic (C18:1Δ9) and linoleic acid (C18:2Δ9,12).^23^ Interestingly, neither stearic acid (C18:0) nor glycerophosphocholine (GPC), the other product of LPC hydrolysis, were found to have antibacterial effects on Gram-positive bacteria.^21,25^

On the other hand, bacteria encode several systems to adapt to the presence of exogenous fatty acids.^26^ For example, Gram-positive bacteria, including *S. aureus*, *E. faecalis*, and pathogenic streptococci, all encode a two-component fatty acid kinase (FakAB) for the uptake of exogenous fatty acids.^27–30^ Whether such a system can confer tolerance and/or resistance to FFAs at physiologically relevant concentrations still awaits validation. Additionally, *S. aureus* encodes the resistance-nodulation-division superfamily efflux pump FarE, which is an important mediator of resistance to antimicrobial fatty acids, including linoleic and arachidonic acids,^31^ with an as yet undefined exportome.

Phospholipases hydrolyze LPC. Based on the cleavage site, phospholipases are grouped into four major types (Fig 1A)^32^: both phospholipase A (PLA) and phospholipase B (PLB) are acyl hydrolases that hydrolyze the ester bond linking the acyl chain with the glycerol. Specifically, PLA_1_ cleaves at the sn-1 position, PLA_2_ cleaves at the sn-2 position, and PLB can cleave at both sn-1 and sn-2 sites. It is worth noting that though some PLA_1_ and PLA_2_ can hydrolyze diacyl-plasmalogens,^33,34^ currently only lysoplasmalogenase is known to hydrolyze the vinyl-ester bond in lysoplasmalogens.^35^ Phospholipase C (PLC) and phospholipase D (PLD) are both phosphodiesterases. PLC cleaves the glycerophosphate bond, releasing a glycerolipid and a phosphate head group, such as the phosphocholine of LPC, while PLD removes the head group, leaving a glycerophosphate lipid. Both humans and bacteria produce phospholipases. Human PLA_2_ has antibacterial activities potentially via direct digestion of bacterial membrane phospholipids.^36^ Bacteria secrete phospholipases as virulence factors for tissue invasion, hemolysis, and phagocytosis evasion.^32^

**Fig 1:**
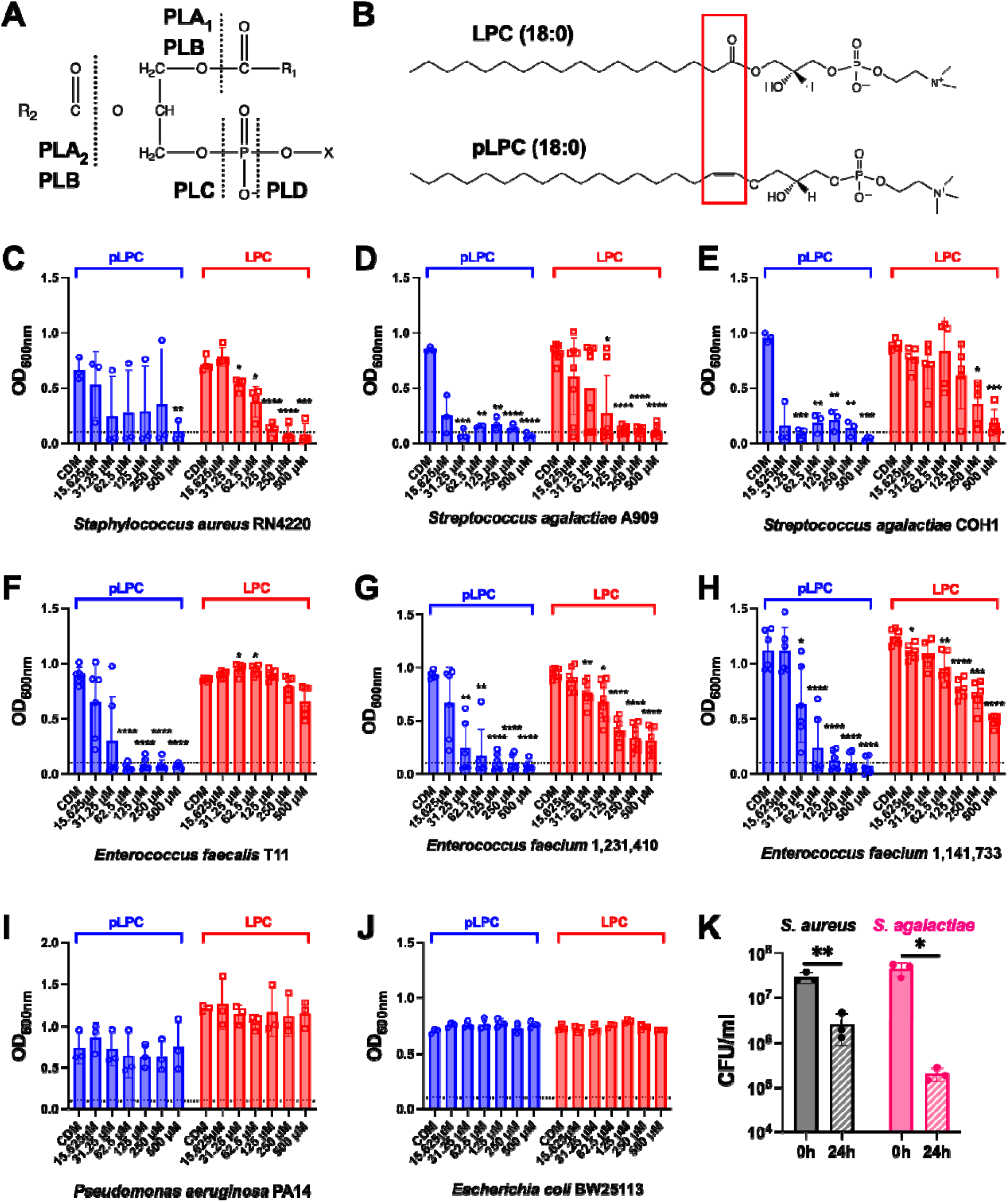
Susceptibilities to lysophosphatidylcholine (LPC) (18:0) and plasmenyl-LPC (pLPC) of selected medically relevant bacterial species. A. Diagram of the cutting sites of phospholipases, including phospholipase A1 (PLA_1_), phospholipase A2 (PLA_2_), phospholipase B (PLB), phospholipase C (PLC), and phospholipase D (PLD). B. Diagrams of the chemical structures of LPC (18:0) and pLPC (18:0). C-J: Plots of OD_600nm_ values for bacteria grown for 24 hours in chemically defined medium (CDM) supplemented with the indicated concentration of LPC (18:0). The dash line indicates the starting OD_600nm_ value (0.1) for the incubation. K: Viable cell counts expressed as CFU/mL for *S. aureus* RN4220 and *S. agalactiae* A909 before (0 h) and after 24 h exposure to 250 μM LPC (18:0). For all panels, ≥ 3 biologically independent replicates were obtained for each strain/condition, with each replicate represented by a data point. Statistical analyses for panels C-J were performed with one-way ANOVA followed by Dunnett’s test for multiple comparisons against the CDM condition. Statistical analyses for panel K were performed with two-tailed T-test. Significant P-values (<0.05) were indicated as: *, 0.05 > *P*-value ≥ 10^-2^; **, 10^-2^ > *P*-value ≥ 10^-3^; ***, 10^-3^ > *P*-value ≥ 10^-4^; ****, 10^-4^> *P*-value.

On the other hand, some bacteria uptake host lipid metabolites for use as substrates for their own lipid biosynthesis. The effects of this lipid scavenging on bacterial pathogenesis and antimicrobial susceptibility *in vivo* remain poorly understood. For example, pathogenic streptococci, *Escherichia coli,* and the plant pathogen *Xanthomonas campestris* all harvest exogenous LPC for the synthesis of PC.^37–40^ Additionally, more than half of *Firmicutes* (recently renamed to *Bacillota*) encode predicted plasmalogen synthesis genes.^41^ These genes mediate plasmalogen production under anaerobic conditions.^41^ The biological functions of plasmalogen lipids in bacteria and the roles of these lipids in host interactions and pathogenesis remain unclear.

Overall, the prior work suggests a link between LPC and pLPC metabolite levels and the risk of infection in an individual. Yet, direct evidence demonstrating the protective roles of these lipid metabolites in the human host is still scarce, let alone a detailed understanding of their antimicrobial mechanisms. In this study, we tested the susceptibility to LPC (18:0) and plasmenyl-LPC (pLPC) (18:0) in selected Gram-negative (*Escherichia coli* and *Pseudomonas aeruginosa*) and Gram-positive (*S. aureus*, *Streptococcus agalactiae*, *E. faecalis*, and *Enterococcus faecium*) pathobionts. Our results demonstrate the potent antimicrobial activity of pLPC against the Gram-positive pathobionts. Interestingly, LPC has pathogen-specific effects, ranging from antimicrobial activity (to *S. agalactiae* and *E. faecium*), to the emergence of spontaneous LPC resistance (in *S. aureus*), to probable inactivation of LPC via lipases and protection of bystander LPC-sensitive pathogens (*E. faecalis*). Overall, our work identifies distinct mechanisms of bacterial resistance to host-derived lipid metabolites with antimicrobial activities.

## Results

### Gram-positive bacteria are sensitive to plasmenyl-lysophosphatidylcholine and have different susceptibilities to lysophosphatidylcholine

The effects of the lipid metabolites LPC and pLPC on bacterial growth were tested at physiologically relevant concentrations. Specifically, in healthy individuals, plasma LPC concentrations range from 125 to 134 μM;^8,42^ yet in individuals with altered health status, such as infections, diabetes, neurodegenerative diseases, and various cancers, the plasma LPC levels fluctuate widely (∼50 to > 300 μM) ^11,43–45^. Thus, a concentration range of 15.625 to 500 μM of LPC was tested. Plasmenyl-LPC was tested with the same concentration range for comparison.

As shown in Fig 1, the growth of Gram-negative bacteria *P. aeruginosa* and *E. coli* was not inhibited by the presence of either LPC or pLPC at all tested concentrations. In contrast, all tested Gram-positive bacteria were susceptible to pLPC with various minimum inhibitory concentrations (MICs). The growth of *S. aureus* and *S. agalactiae* was completely inhibited by LPC (Fig 1C-D). A significant decrease in their viable cell numbers after 24-hour exposure to 250 μM LPC (Fig 1K) indicates a bactericidal effect of LPC. Both *E. faecalis* and *E. faecium* could grow with the presence of LPC (Fig 1F-H). Significantly higher growth of *E. faecalis* was even observed when it was incubated with LPC at 31.25 and 62.5 μM, suggesting that *E. faecalis* is resistant to LPC.

### Enterococcus faecalis deactivates extracellular lysophosphatidylcholine

pLPC only differs from LPC at the bond linking the acyl chain to glycerol (Fig. 1B), which changes its susceptibility to acyl hydrolase. It is possible that *E. faecalis* deactivates LPC by a lipase(s) that is ineffective in hydrolyzing pLPC. Indeed, LPC is absent in the total lipids harvested from *E. faecalis* incubated with 50 μM LPC (Fig S1). As a comparison, LPC is still present in the total lipids extracted from *S. aureus* grown with 50 μM LPC (Fig S1). Additionally, neither *E. faecalis* nor *S. aureus* were found to produce PC in the presence of LPC (Fig S1). PC is produced by pathogenic and commensal streptococci by scavenging exogenous LPC.^37,38^ Such results suggest that *S. aureus* and *E. faecalis* lack this pathway.

To verify the extracellular deactivation of LPC by *E. faecalis*, LPC-containing media were pulse-treated by *E. faecalis*, filtered, and then used to grow LPC-susceptible bacteria. Specifically, mid-log phase *E. faecalis* were added to fresh chemically defined medium (CDM) containing 250 μM LPC to a final concentration at 10^9^ CFU/mL, followed by a 1-hour incubation at 37°C, after which the bacterial cells were removed by centrifugation and filtration (Fig 2A, Set 1).

**Fig 2:**
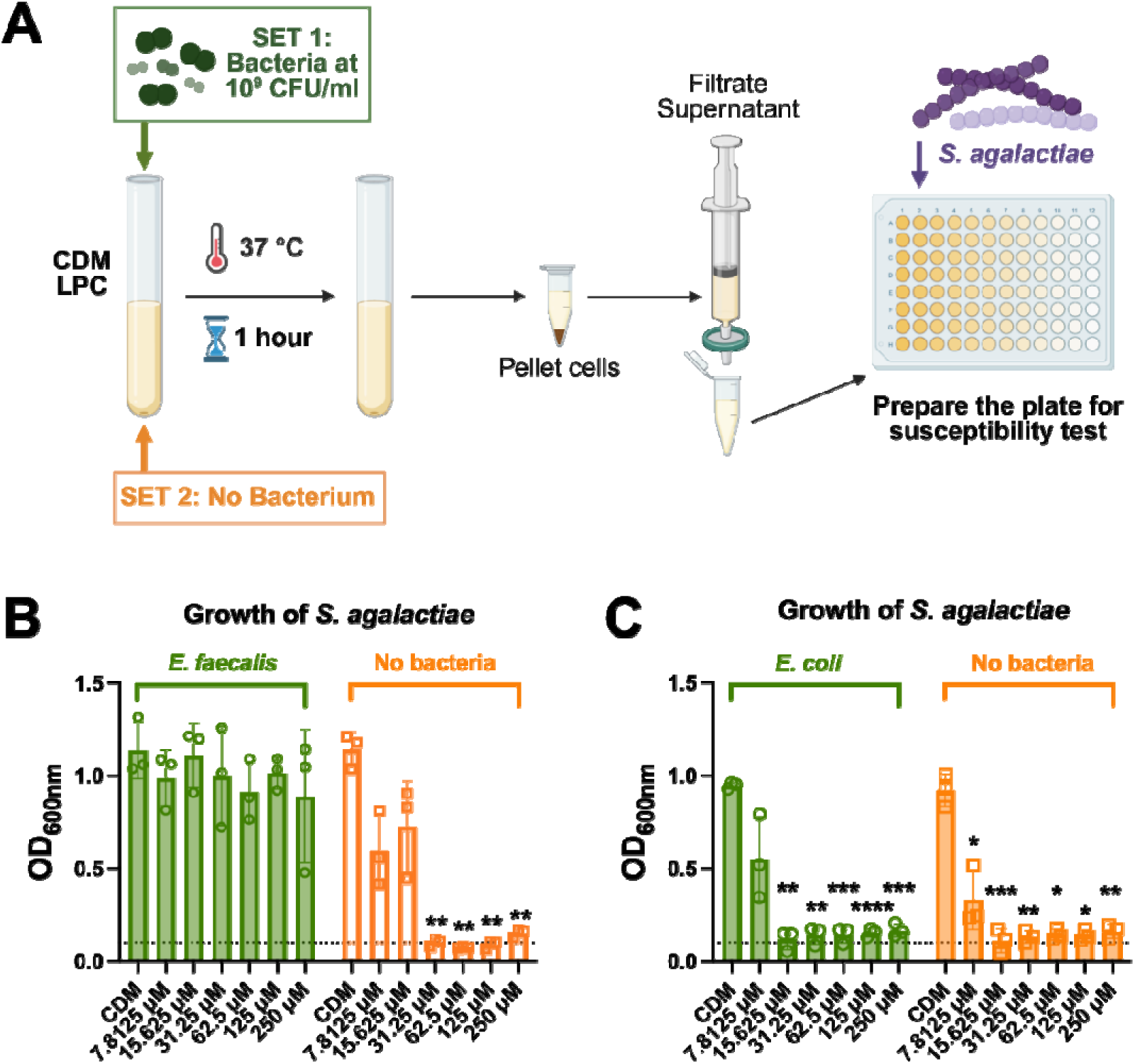
***E. faecalis*, but not *E. coli*, deactivates extracellular LPC.** Panel A: Experimental procedure diagram. Panels B and C: Plot of the OD_600nm_ values of *S. agalactiae* A909 grown overnight in CDM supplemented with LPC (18:0), with or without *E. faecalis* (panel B) or *E. coli* (panel C) pre-treatment. Specifically, CDM containing 250 μM LPC (18:0) was pulse-treated for 1 h by 10^9^ CFU/mL of either *E. faecalis* or *E. coli* (Set 1 in Panel A) or was left untreated but processed with the same procedures (Set 2 in Panel A). The indicated LPC concentrations were calculated based on the pre-treatment LPC concentrations. Dashed lines indicate the starting OD_600nm_ value (0.1) of the *S. agalactiae* cultures. For each tested condition, at least 3 biologically independent replicates were obtained, with each replicate represented by a data point. Statistical analyses were performed with one-way ANOVA followed by Dunnett’s test for multiple comparisons against the CDM condition. Significant *P*-values (<0.05) were indicated as: **, 10^-2^ > *P*-value ≥ 10^-3^; ***, 10^-3^ > *P*-value ≥ 10^-4^; ****, 10^-4^ > *P*-value.

Bacterial free LPC-containing CDM was incubated, centrifuged, and filtered at the same time (Fig 2A Set 2), thus controlling for any potential loss of LPC due to the experimental procedures. LPC-sensitive *S. agalactiae* could grow in *E. faecalis* pre-treated media but not in the control media (Fig 2B), indicating a successful deactivation of the LPC-mediated antibacterial activity by *E. faecalis*. As a comparison, media pulse-treated by *E. coli* in the same manner still maintained its ability to limit growth of LPC-sensitive *S. agalactiae* (Fig 2C).

### Spontaneous LPC-resistant S. aureus mutants lack odd-chain fatty acids and have induced expression of the efflux pump FarE

After the initial measurement of bacterial growth (after 24 hours of incubation) in the presence of LPC (Fig 1), the plates were incubated on the benchtop and observed regularly for turbidity changes. Within four days, the LPC-containing cultures of both *S. aureus* and *S. agalactiae* frequently became turbid. Stable LPC-resistant mutants could only be obtained from *S. aureus*, but not *S. agalactiae*, as only *S. aureus* grew after re-inoculation into fresh CDM with LPC. Specifically, independent spontaneous LPC-resistant mutants were obtained from the *S. aureus* laboratory strain RN4220 (3 mutants: B3, B4, B6), methicillin-sensitive clinical isolate MSSA476 (R1, R2, R3), and methicillin-resistant clinical isolate USA300 (R2, R4) (Table 1).

**Table 1:** Mutations in *S. aureus* lysophosphatidylcholine-resistant strains from this study.

| **Gene** | **Strains and Mutants** |  | **Function** | **Reported antimicrobial resistance in *S. aureus*** |
|---|---|---|---|---|
| *farR* | RN4220 | B3: Val115Ile | Regulator of fatty acid efflux pump (FarE) and other metabolic and virulence associated genes.<sup>46</sup> | Spontaneous FarR<sub>Asp94Tyr</sub> and FarR<sub>Val115Ile</sub> mutants are resistant to lipid-based antimicrobials.<sup>46,49</sup> |
|  |  | B6: Asp94Tyr |  |  |
|  | USA300 | R2: Glu151Lys |  |  |
|  |  | R4: Val115Ile |  |  |
| *fabF* | RN4220 | B4: Phe230Ile | Fatty acid biosynthesis elongation condensing enzyme.<sup>47</sup> | Spontaneous FabF<sub>Pro137Leu</sub> mutation contributes to daptomycin resistance.<sup>51</sup> |
|  | MSSA476 | R1: Ser397Ile |  |  |
| *walK* | MSSA476 | R2: Gln371del | Sensor kinase of a two-component system that controls cell wall metabolism.<sup>48</sup> | Spontaneous mutations lead to co-resistance to vancomycin and daptomycin.<sup>48</sup> |
|  |  | R3: Ser9Pro |  |  |

Genome resequencing analyses were performed to identify unique single-nucleotide polymorphisms (SNPs) that arose in these LPC-resistant isolates compared to their isogenic wild type (WT) parental strain. Mutations in three chromosomal genes were repeatedly detected (Table 1): *farR*, encoding a TetR family transcriptional regulator that controls multiple cellular functions;^46^ *fabF*, encoding a condensation enzyme that elongates fatty acids;^47^ and *walK* (or *yycG*), encoding the sensor kinase of a two-component system that is essential for viability.^48^ Spontaneous mutations in all three genes were previously observed in *S. aureus* clinical isolates and contribute to multidrug resistance.^31,46,48–51^

Considering that all three mutated genes are linked with lipid metabolism, lipidomic analyses were performed. A dramatic decrease in odd-chain fatty acids was observed in all of the tested LPC-resistant mutants (Table 2, Fig S2-4). Previous studies indicate that FarR mutations can lead to the overexpression of the lipid efflux pump FarE.^31,46^ Indeed, qRT-PCR confirmed a significant increase of *farE* transcripts in RN4220 *farR* mutants (Fig 3A), B3 (FarR_Val115Ile_) and B6 (FarR_Asp94Tyr_), as well as the *fabF* mutant B4 (FabF_Phe230Ile_).

**Fig 3:**
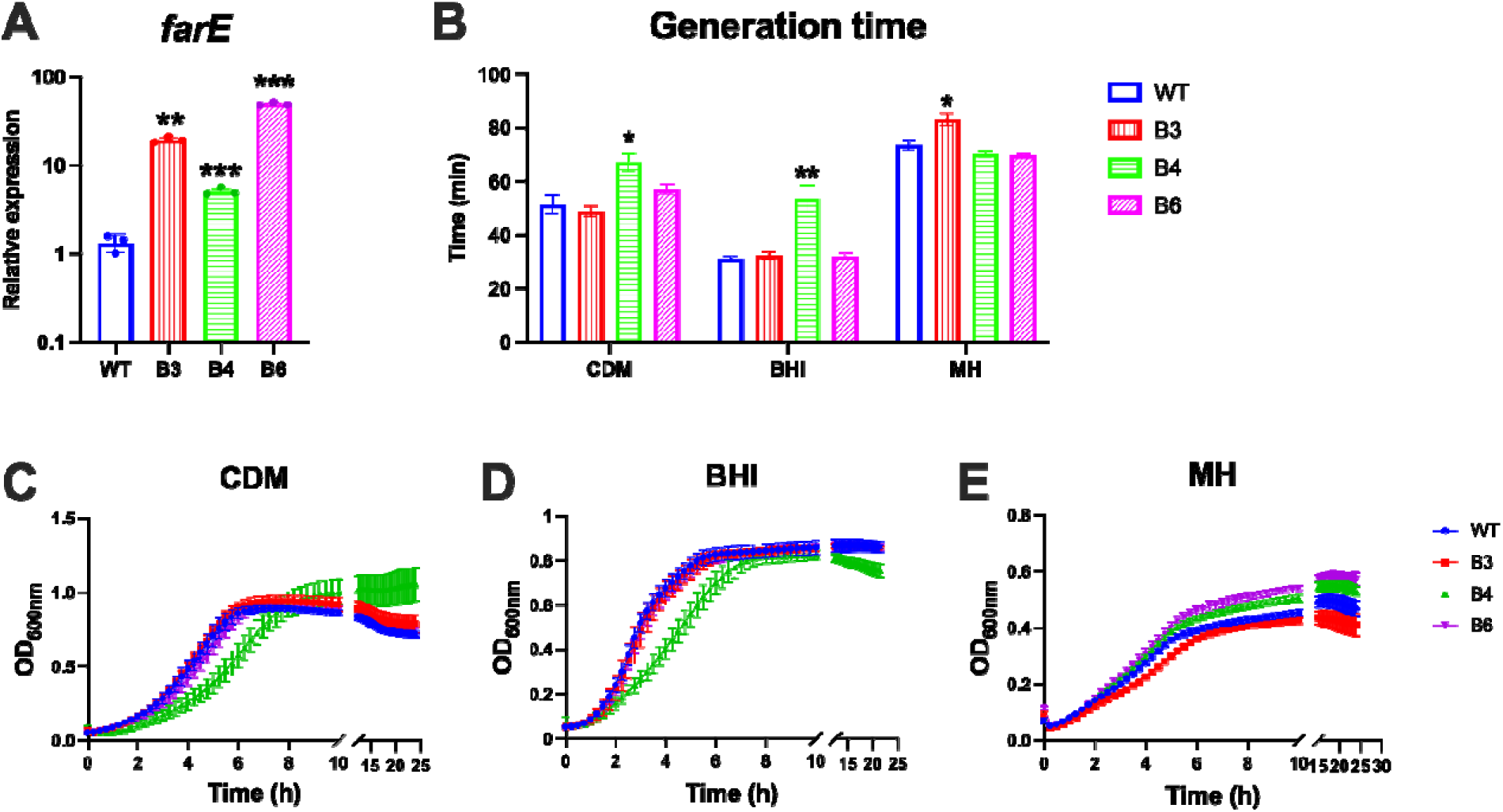
***S. aureus* RN4220 LPC-resistant mutants have elevated *farE* transcript levels and altered growth relative to the wildtype (WT).** Panel A. The relative expression levels of *farE* transcripts measured with qRT-PCR and calculated with the ΔΔCt method normalizing to the levels of 16S rRNA (*rplD*). Panel B. Generation time of *S. aureus* WT and mutant strains grown in the indicated media. Panels C-E. Growth curve for *S. aureus* WT and mutant strains grown at 37°C in chemically defined medium (CDM, panel C), brain heart infusion medium (BHI, panel D), and Mueller Hinton medium (MH, panel E). Values of OD_600nm_ were measured at 15 min intervals for 24 hours. Color coding for all panels: blue, WT; red, mutant B3 (FarR_Val115Ile_), green, B4 (FabF_Phe230Ile_), and purple, B6 (FarR_Asp94Tyr_). For each tested condition, at least 3 biologically independent replicates were obtained. Statistical analyses were performed with one-way ANOVA followed by Dunnett’s test for multiple comparisons against data of the WT samples. Significant *P*-values (<0.05) were indicated as: *, 0.05 > *P*-value ≥ 10^-2^; **, 10^-2^ > *P*-value ≥ 10^-3^; ***, 10^-3^ > *P*-value ≥ 10^-4^.

**Table 2:** Fatty acids detected in *S. aureus* wild-type strains and their spontaneous LPC-resistant mutants

| **Strain** |  | **C14:0** | **C15:0** | **C16:0** | **C17:0** | **C18:0** | **C19:0** | **C20:0** | **C21:0** |
|---|---|---|---|---|---|---|---|---|---|
| RN4220 | WT | Yes | Yes | Yes | Yes | Yes | - | Yes | Yes |
|  | B3 | - | - | - | - | Yes | - | Yes | - |
|  | B4 | Yes | - | Yes | - | Yes | - | Yes | - |
|  | B6 | Yes | - | Yes | - | Yes | - | Yes | - |
| MSSA476 | WT | - | - | Yes | Yes | Yes | Yes | Yes | Yes |
|  | R1 | - | - | Yes | - | Yes | - | Yes | - |
|  | R2 | - | - |  | - | Yes | - | - | - |
|  | R3 | - | - |  | - | Yes | - | Yes | - |
| USA300 | WT | - | Yes | Yes | Yes | Yes | Yes | Yes | Yes |
|  | R2 | - | - | Yes | - | Yes | - | Yes | - |
|  | R4 | - | - |  | - | Yes | - | Yes | - |

### LPC-resistant S. aureus strains have altered antibiotic susceptibilities and dampened LPC-induced stress responses

Previous studies reported pleiotropic phenotypes in *S. aureus* with a point mutation in *farR* (FarR_Cys116Arg_), leading to increased pathogenicity and multi-drug resistance compared to the WT strain.^46,49^ Among the three FarR variants reported in this study, the FarR_Val115Ile_ was detected in two strains among >2,000 isolates of *S. aureus* from various sources (Fig 4) and in strains of the clonal complex 5 hospital-associated methicillin-resistant *S. aureus*,^52^ suggesting its relevance to adaptations to host environments.

**Fig 4:**
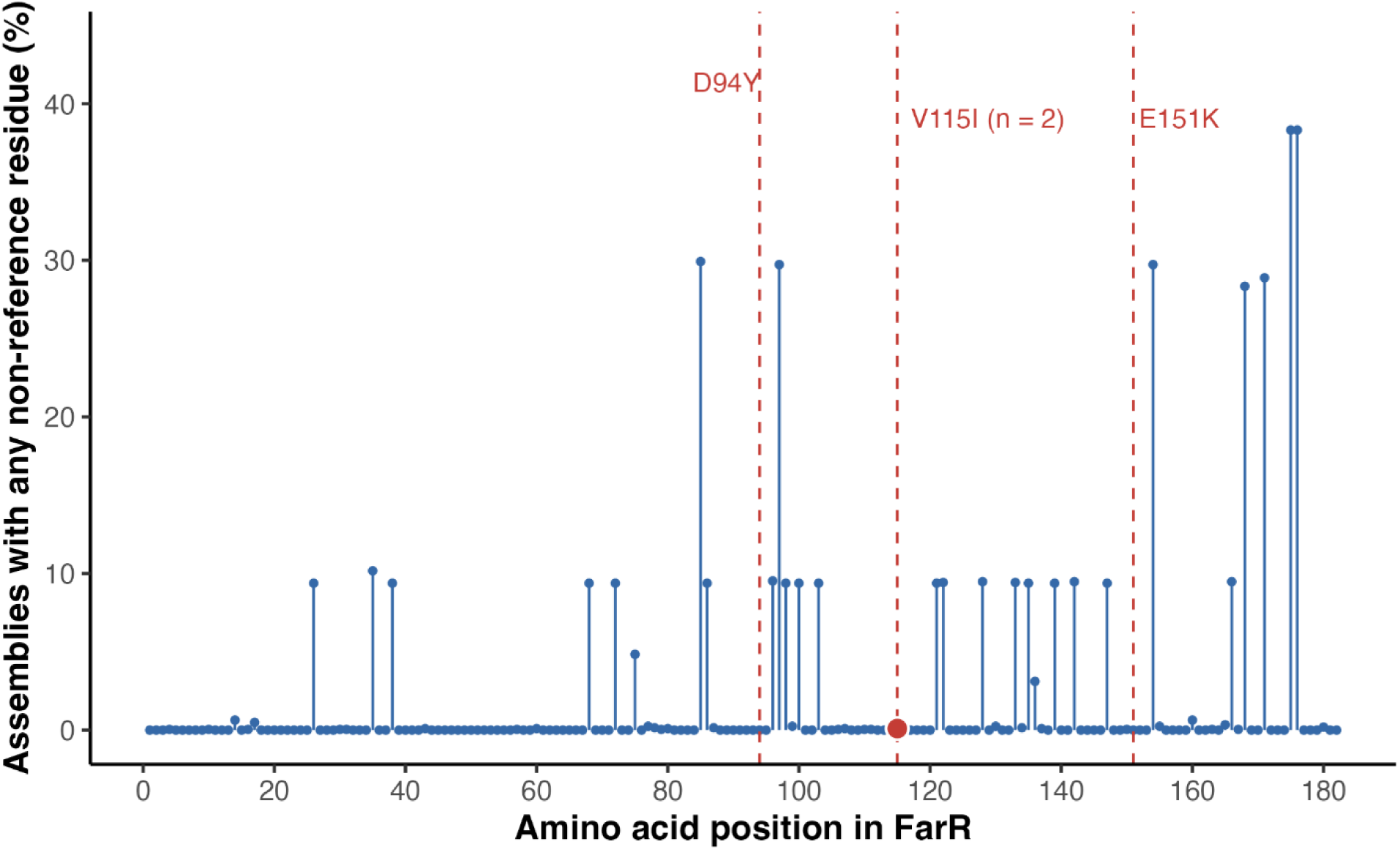
FarR variants identified from publicly available *S. aureus* genome sequences and this project. Sites of polymorphism across 2025 FarR amino acid sequences identified from analysis of 2042 complete *S. aureus* Refseq assemblies (amino acid position along the X axis). Polymorphism sites are marked by blue solid lines and dots. The three substitutions identified in this work are indicated by red dashed line - D94Y, V115I, and E151K. Only V115I was detected in this cohort, occurring in two assemblies: GCF_003186105.1 of *S. aureus* MOK063 and GCF_041729565.1 of *S. aureus* 23EV612; per genome metadata, both of these strains were isolated from bovine milk, from Ireland and New Zealand, respectively.

Both phenotypic and transcriptomic analyses were performed to explore the effects of FarR variations. Specifically, compared to the *S. aureus* RN4220 WT strain, mutant B4 (FabF_Phe230Ile_) has significant growth deficiency in CDM and BHI (Fig 3); and mutant B3 (FarR_Val115Ile_) has significant growth deficiency in Mueller Hinton (MH) medium, which is the standard medium used for antimicrobial susceptibility tests.^53^ Mutant B3 (FarR_Val115Ile_) demonstrated decreased susceptibility to daptomycin (Table 3), and the same trend was also observed in B6 (FarR_Asp94Tyr_); while mutant B4 (FabF_Phe230Ile_) had decreased susceptibility to fosfomycin. Additionally, as indicated in Table 3, both B3 and B4 have increased susceptibility to gentamicin, and all tested mutants have a slight increase in their susceptibility to vancomycin.

**Table 3:** Minimum inhibition concentrations (μg/ml) of selected antibiotics in *S. aureus* RN4220.

| **Strain** | **Antibiotic<sup>a</sup>** |  |  |  |
|---|---|---|---|---|
|  | Daptomycin | Vancomycin | Gentamicin | Fosfomycin |
| Wild type | 1 (0.75 – 2) | 2 (2) | 0.25 (0.25) | 0.19 (0.125 – 0.19) |
| B3 | 1.5 (1 – 1.5) | 1.5 (1.5 – 2) | 0.125 (0.125) | 0.125 (0.125) |
| B4 | 1 (1) | 1.5 (1.5) | 0.094 (0.064 – 0.125) | 0.5 (0.38 – 0.5) |
| B6 | 1.5 (1.5 – 2) | 1.5 (1.5 – 2) | 0.25 (0.25) | 0.19 (0.125 – 0.19) |

<sup>a</sup>Measurement of the minimum inhibitory concentration was conducted with E-test strips (N=3). The median MIC is shown, with the range of MICs detected in parentheses.

Transcriptomic analyses were performed to compare responses to LPC in the *S. aureus* RN4220 WT and B3 (FarR_Val115Ile_) mutant. Total RNA was harvested from mid-log phase cells that were incubated at 37°C for 1 hour with and without the presence of 50 μM LPC. When LPC was absent, only *farRE* were differentially expressed between WT and B3, with significantly increased expression levels observed in B3 (*farR* > 4-fold; *farE* > 106-fold, compared to WT; RPKM values demonstrated in Fig 5A), indicating that such point mutation only directly modulates *farRE* expression levels in a non-stressed condition. When LPC is present, a total of 111 genes were differentially expressed between WT and B3 (Fig 5B), with *farE* and *farR* still being the highest up-regulated genes in B3 (*farR* > 5-fold; *farE* > 30-fold). Other differentially regulated genes include those involved in stress responses (*groES*, *groL*, and *clpB*, upregulated in WT), histidine biosynthesis (*his* genes, upregulated in WT), purine biosynthesis (*pur* genes, downregulated in WT), iron-regulated surface determinants (*isd* genes, down regulated in WT), and staphyloferrion B biosynthetic genes (*sbnGHI*, downregulated in WT).

**Fig 5:**
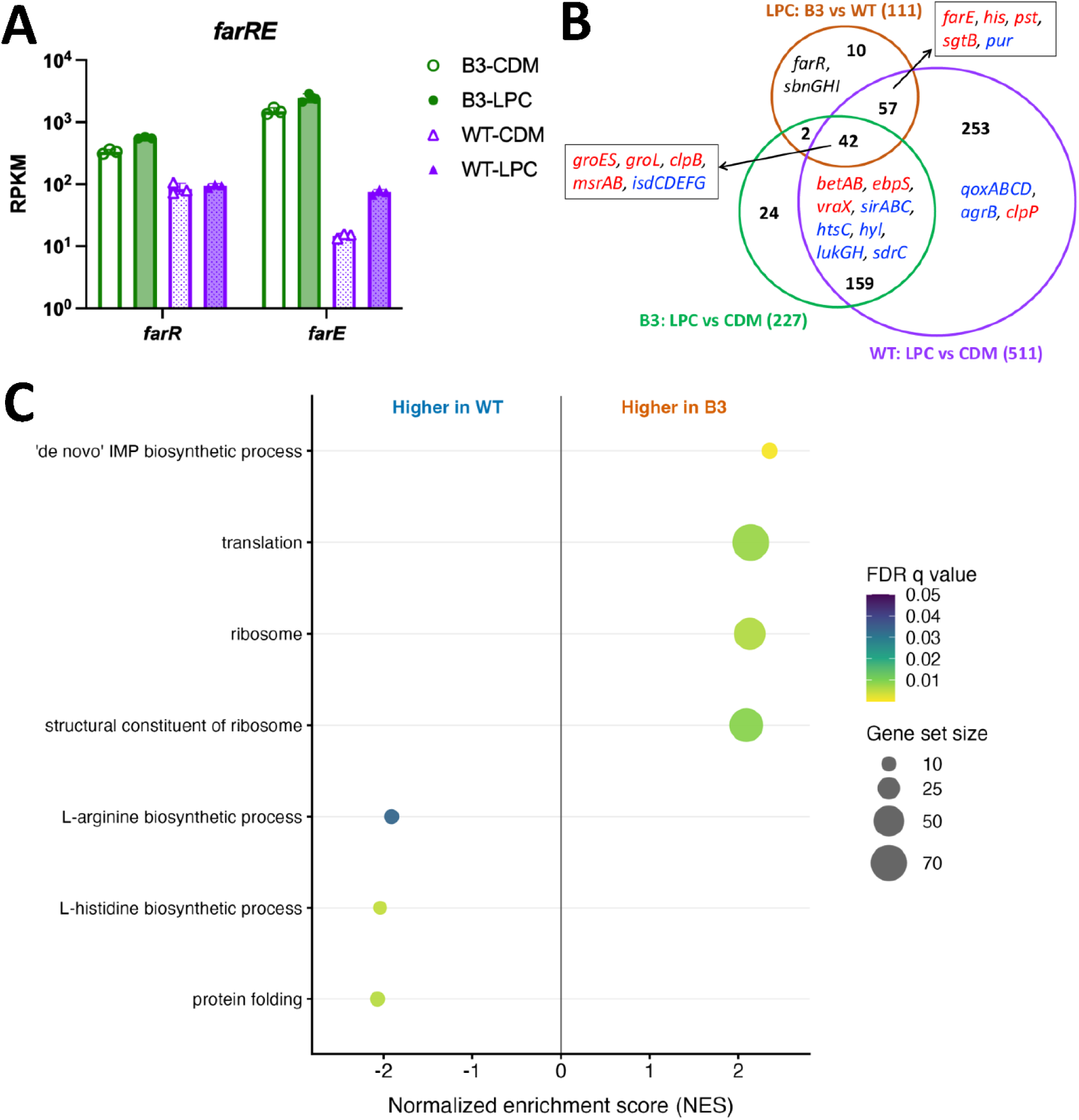
**Differentially expressed genes in *S. aureus* WT and B3 with and without the exposure to LPC (18:0)**. Total RNAs were harvested from WT and B3 strains grown to mid-log phase followed with the addition of either LPC (18:0) to 50 μM (the LPC condition, indicated by the solid symbols and plot bars in panel A) or an equal volume of methanol (the CDM condition, indicated by the empty symbols and corresponding plot bars in panel A). Three biologically independent replicates were obtained for each sample at each tested condition. Panel A. RPKMs of *farR* and *farE*. Panel B. Venn Diagram of the differentially expressed gene profiles, including the WT genes differentially expressed with and without LPC (purple, WT: LPC vs CDM), B3 genes differentially expressed with and without LPC (green, B3: LPC vs CDM), and genes differentially expressed between B3 and WT when both strains were exposed to LPC (orange, LPC: B3 vs WT). Gene examples were labeled within each corresponding section, except for the ones of the 42 and 57 shared genes, which were indicated in the arrowed boxes. Genes upregulated by LPC are indicated with red fonts; genes downregulated by LPC are indicated with blue fonts. Gene *farR* and *sbnGHI* are upregulated in B3 compared to WT. Panel C. Gene ontology (GO) enrichment analytic results comparing B3 and WT with LPC exposure. Genes were ranked by the DESeq2 Wald statistic, and all GO terms with a GSEA FDR q value < 0.05 are shown. Positive normalized enrichment scores (NES) indicate enrichment in B3, whereas negative scores indicate enrichment in WT. Dot size represents the gene-set size (number of ranked genes assigned to each GO term) and dot color represents the Benjamini-Hochberg adjusted p-value.

Compared to no LPC conditions, LPC stimulated differential expression of 511 genes in WT and 227 genes in B3 (Fig 5B). Between WT and B3, 201 genes were shared, none of which were regulated in the opposite direction. Generally, in both WT and B3, LPC stimulated an upregulation of stress response genes, including *groL*, *groES*, *clpB*, and *msrAB*, and a downregulation of iron utilization genes, such as the *isd* genes and the iron-regulated sortase gene *srtB*. Virulence genes also appeared in the 201 shared genes, among which genes encoding hyaluronidase (*hyl*), leukocidin (*luk*), and fibrinogen-binding protein (*sdrC*) were down-regulated by LPC, and genes encoding the elastin-binding protein (*ebpS*) and a secreted complement C1q inhibitor (*vraX*) were up-regulated by LPC.

Genes that were uniquely differentially expressed in B3 with LPC exposure include the enterotoxin-like toxin *selX* (down-regulated by LPC), extracellular adherence protein *eap* (down-regulated by LPC), and the gamma-hemolysin HlgAB subunit A *hlgA* (up-regulated by LPC).

Genes that were only found to be differentially expressed in WT include the ones encoding the protein essential for quorum sensing peptide biosynthesis (AgrB),^54^ ribosomal proteins, and quinol oxidase (QoxABC),^55^ all of which were downregulated by LPC; while genes encoding the metabolic and virulence regulator CodY,^56^ lipoteichoic acid synthase LtaS,^57^ and cell membrane stress sensing accessory protein LiaF,^58^ as well as those involved in riboflavin biosynthesis, were all up-regulated by LPC.

Gene ontology enrichment analyses (GSEA) comparing B3 and WT in response to LPC exposure found the following biological processes to be mostly affected (FDR q< 0.05; Fig 5C and S5): 1) Terms enriched in WT included histidine biosynthesis (including *hisABDFGH*), protein folding stress responses (including *groL*, *clpB*, *dnaK*, and *dnaJ*) and L-arginine biosynthesis. 2) Terms enriched in B3 included purine biosynthesis (including the *pur* genes), ribosomal subunits and translation. Among processes highlighted by the gene-level analysis, response to metal ions (NES=1.84, nominal P < 0.001, FDR q=0.073) and riboflavin biosynthesis (NES=1.71, nominal P=0.0037, FDR q=0.194) had WT-directed enrichment scores but did not meet the FDR threshold.

Overall, our transcriptome studies identify a broad transcriptional response of WT *S. aureus* to LPC exposure, likely due to the antimicrobial effects of this lipid on the *S. aureus* cell. The LPC-resistant strain mounted a fraction (∼39%) of this response. Notably, the LPC-resistant strain was not “blind” to LPC and still mounted a transcriptional response to LPC, with some unique responses relative to the wild-type. Whether these responses are due to a continued antimicrobial effect of LPC against this strain or perhaps to direct sensing of LPC (speculatively) as a host-derived signal remains to be determined.

## Discussion

In this study, we demonstrated diverse susceptibility profiles of human-associated bacteria to the human lipid metabolites LPC (18:0) and pLPC (18:0) at physiologically relevant concentrations.

Gram-negative bacteria, including *E. coli* and *P. aeruginosa*, were not susceptible to either LPC or pLPC. In contrast, all tested Gram-positive bacteria were susceptible to pLPC. Such antibacterial property of pLPC suggests a potential role of this lipid in preventing or mitigating infections, particularly in the brain and heart where it is abundant, which awaits further experimental verification. While LPC can effectively kill *S. aureus* and *S. agalactiae*, it does not completely inhibit the growth of enterococci; instead, it even promotes the growth of *E. faecalis* at low concentrations. Additionally, *E. faecalis* can metabolize extracellular LPC, indicated by the absence of LPC in lipidomic analyses as well as the growth of *S. agalactiae* in *E. faecalis*-treated LPC-containing media. Spontaneous LPC-resistant *S. aureus* arose *in vitro*, and the strains all demonstrated altered lipid profiles, with several isolates containing point mutation in *farR* that leads to increased expression of the efflux pump FarE, which possibly contributed to the observed resistance. Collectively, these results indicate distinct mechanisms of bacterial resistance to host-derived lipid metabolites with antimicrobial activities.

Gram-positive bacteria, including *S. aureus*, pathogenic streptococci, and enterococci, are all known to possess the exogenous fatty acid uptake system FakAB, which has been implicated in protection against antimicrobial lipids.^26,27,30,59^ Additionally, streptococci scavenge environmental LPC for PC production, a potential function of which in resisting antimicrobial lipids is also unknown.^37,38^ Our susceptibility data suggest that neither the Fak nor the LPC-incorporating pathway is involved in protecting the Gram-positive bacteria tested here against LPC at physiologically relevant concentrations, as both *S. aureus* and *S. agalactiae* are susceptible. Instead, *E. faecalis*, which cannot produce PC from utilizing LPC, potentially resist LPC by lipase(s), as indicated by its susceptibility to pLPC and effective extracellular deactivation of LPC. So far, no lipase, and particularly no acyl hydrolase, has been experimentally characterized in *E. faecalis*. Identification of the lipase(s) responsible for mediating the observed LPC-resistance can provide novel target(s) for designing antimicrobials that specifically limit bacterial growth under host-associated conditions.

Meanwhile, we report spontaneous LPC-resistant *S. aureus* strains with point mutations in genes involved in lipid metabolism. Spontaneous mutations of these genes have been observed in clinical isolates with increased multidrug resistance. Specifically, *fabF* point mutations lead to daptomycin resistance in both *S. aureus* and *E. faecium*.^51,60^ Point mutations in *S. aureus walKR* lead to intermediate vancomycin resistance and daptomycin nonsusceptibility.^48^ Point mutations in *S. aureus farR* have been frequently reported in both laboratory lipid antimicrobial adaptation studies and multidrug resistant clinical isolates,^49,50,52^ particularly in the clinically prominent MRSA clonal complex 5.^52^ All these studies highlight the importance of lipid metabolism in bacterial pathogenesis. Increased *farE* expression is observed in both *fabF* and *farR* mutants, suggesting a potential crosstalk between these metabolic pathways. The shared phenotype of dramatically decreased production of odd-chain fatty acids in these mutants suggests that these three pathways collaboratively shape the bacterial lipid composition. RNA-seq results indicate an alleviation of LPC-induced stress in the FarR_Val115Ile_ mutant. Interestingly, LPC induced a downregulation of iron utilization genes in both the WT and mutant, and a few virulence genes were differentially regulated between WT and mutant, indicating an altered virulence status of the FarR_Val115Ile_ mutant. Future analyses are needed to confirm such transcriptomic changes in clinical isolates and reveal the detailed lipid metabolic regulatory networks in *S. aureus*. A comprehensive understanding of lipid-mediated host-*S. aureus* interactions is required to design novel therapeutics targeting this axis of *S. aureus* pathogenicity.

## Materials and Methods

### Bacterial strains and growth conditions

Unless indicated otherwise, all bacteria were grown at 37°C. For routine culture, Gram-negative bacteria were grown in Luria-Bertani (LB) medium, *S. aureus* and enterococci were grown in Brain-Heart Infusion (BHI) medium, and streptococci were grown in Todd-Hewitt (TH) medium. Chemically defined medium (CDM) was prepared as previously described.^61^

### Antibiotic susceptibility testing

Antibiotic susceptibility tests were performed using E-test strips (BioMet) per the manufacturer’s instructions. Specifically, a single colony was inoculated into Mueller-Hinton (MH, BD Biosciences) broth over-night followed by spreading onto MH agar plates. Daptomycin MICs were measured through cultivation with MH media supplemented with 50 mg/L calcium as instructed by EUCAST.^53^

### LPC susceptibility assays and isolation of spontaneous LPC-resistant mutants

The lipids used in this study were LPC (18:0) (Avanti Polar Lipids, SKU 855775), and C18 (Plasm) LPC (Avanti Polar Lipids, SKU 852465P) (pLPC). Stock solutions at 250 mM were made with methanol. For all tested bacteria, no significant growth inhibition was observed when methanol was added to a final concentration of 0.2% (v/v) (Fig S6).

The minimum inhibitory concentration (MIC) of LPC for each tested bacterium was measured in a serial microdilution method modified from EUCAST guidelines.^53^ Specifically, CDM was used instead of MH medium to enable control of the concentrations of the added lipid metabolites.

The LPC stock solution was diluted to 1 mM in CDM as the working solution. After adding 200 μL working solution into all row C wells of a 96-well plate, 100 μL was transferred into wells in row D to mix with 100 μL CDM. Such dilution was performed serially down the following rows of the plate, leaving 100 μL of LPC dilutions in each well from row C to H. Overnight bacterial cultures grown in CDM were diluted to an OD_600nm_ value of 0.2, 100 μL of which was added to wells from row B to H, yielding 200 μL bacterial culture at OD_600nm_ 0.1 in each well from row C to H. Row B wells served as the positive control for bacterial growth with the addition of 100 μL CDM; row A wells served as the media blank control with the addition of 200 μL CDM. After overnight incubation at 37°C, OD_600nm_ values were measured in a plate reader (Synergy II), after briefly shaking the plate, to quantify bacterial growth. For each bacterium, at least three biological replicates and two technical replicates were measured. Statistical analyses were performed with one-way ANOVA followed by Dunnett’s test for multiple comparisons against the CDM condition. Significance was determined by *P*-value < 0.05.

Two bacterial strains were further assessed for their changes in viable cell number in response to the addition of LPC. Specifically, overnight bacterial cultures were diluted to OD_600nm_ 0.1 in 5 mL pre-warmed CDM with 500 μM LPC. Colony forming units (CFUs) per mL culture were quantified by serial dilution and plating at time 0 (i.e. immediately after the dilution) and after 24 hours of incubation with LPC.

Bacteria observed to have no visible growth in the presence of LPC in the 96-well plate assays were incubated for up to 4 days at room temperature, at which point turbidity was observed for some cultures. Cultures that turned turbid were inoculated into freshly made CDM with LPC at 250 and 500 μM. After overnight incubation, 100 μL of each turbid cultures were streaked onto a BHI agar plate for single-colony isolation. Individual colonies were then inoculated into CDM with 500 μM LPC and incubated overnight. Any resulting turbid cultures were cryopreserved.

### Genome resequencing and analysis

Genomic DNA was extracted with the DNeasy Blood & Tissue Kit (Qiagen) per the manufacturer’s instructions. Illumina genome sequencing was performed by SeqCenter (Pittsburgh, PA). Sequencing results were analyzed with CLC Genomics Workbench (version 23; Qiagen). Briefly, sequence reads were assembled to appropriate reference genomes (NZ_CP101124.1 for *S. aureus* RN4220, NC_007793.1 for *S. aureus* USA300, and NC_002953 for *S. aureus* MSSA476) and variants were detected using default parameters. Non-synonymous variants were considered for further analysis.

### Assessment of bacterial community protection

Community protective effects were assessed with cell-free culture supernatants obtained from cultures that were pulse-treated by lipid-resistant bacteria. Cell-free culture supernatant was obtained with the following procedures (Fig 2A): bacterial cells harvested from overnight cultures grown in CDM were resuspended in 5 mL fresh CDM with 250 μM LPC to an estimated final density of 10^9^ CFU/mL. After incubation at 37°C for 1-hour, bacterial cells were removed by centrifugation at 8,000 g for 10 minutes followed by filtration through a 0.22 μm pore-size sterile filter. Controls without the addition of bacterial cells were prepared with the same concentration of LPC, incubated, centrifuged, and filtered synchronically. The harvested supernatants were used for susceptibility tests as described above.

### Lipid Extraction

Total lipids were extracted with the acidic Bligh-Dyer method and analyzed as described previously.^62^ At least 5 mL cultures were grown in CDM to exponential phase, then LPC was added to 50 μM final concentration, followed by 2 hours incubation at 37°C and lipid extraction. Cultures without the LPC addition but incubated for the same time were extracted as the control. A spike-in control, PC (13:0/13:0), was added to the samples at a final concentration of 100 μM immediately before lipid extraction. See supplementary materials for details of lipidomic analysis.

### RNA extraction, quantitative reverse transcriptase PCR (qRT-PCR), and RNA-sequencing analyses

Overnight bacterial cultures were diluted to an OD_600nm_ value of 0.1 with fresh CDM into two sets and incubated at 37°C. When the OD_600nm_ values were in the range of 0.4-0.5, LPC was added into one set of the culture to a final concentration of 50 μM, and an equal volume of methanol was added to the other set of cultures. After 1 hour incubation, cells were pelleted with centrifugation at 10,000 x g for 10 min. Cell pellets were resuspended with 450 μL CDM and mixed with 900 μL RNAProtect (Qiagen). After incubation at room temperature for 10 minutes, cells were pelleted at 5000 x g for 10 minutes and stored at -80°C prior to further processing.

Total RNA was harvested with the same procedure described previously from *S. aureus*.^63^ DNase treatment of RNA samples and generation of cDNA were conducted as previously described.^63^ Primers used for qRT-PCR were designed via the online program Primer3 with 60±2°C set as the optimized annealing temperature.^64^ Primer pair rplD-F CGTGGCGGTGGTATCGTATT and rplD-R ACTTGAACGCCAGGGATGTT was used for the expression detection of ribosomal gene *rplD*, and primer pair farE-F ACGGAGATTTCGGGACAAAGT and farE-R GCTAGCATTGGTCCTGTCGT was used for the expression detection of *farE*. qRT-PCR reactions were prepared with AzuraQuantTM Green Fast qPCR Mix LoRox (Azura Genomics) per the manufacturer’s instructions, using 1 μL cDNA per reaction for the detection of 16S rRNA and 5 μL cDNA per reaction for the detection of target genes. qRT-PCR reactions were performed on the MyGo Pro machine (Azura Genomics) and analyzed with the MyGo Pro PCR Software (v3.2, Azura Genomics). Relative transcript levels of the target gene *farE* were calculated with the ΔΔCt method with normalization to the Ct numbers of *rplD*.

Illumina sequencing of the total RNA samples was performed by SeqCenter (Pittsburgh, PA). Analyses of the sequencing results were performed with CLC Genomics Workbench (version 23, Qiagen). The *S. aureus* RN4220 genome sequence (NZ_CP101124.1, taxid: 561307) was used as the reference. RPKM values of each annotated gene were calculated and used for the analysis of differential expressions at the transcript level. The validity of replicates was confirmed with principal-component analysis and scatterplot with linear regression using the RPKM values. The R^2^ values of the comparisons between replicates under the same testing condition were all above 0.95 (Table S1). RNA sequencing data for all ORFs are shown in Table S2. Differentially expressed genes were determined by an absolute fold change of > 2.0, a false discovery rate (FDR) P value of < 0.01, and a maximum RPKM group mean of > 10.

### Gene set enrichment analyses (GSEA)

To generate ranked gene lists for GSEA, raw gene counts from all 12 samples were modeled using DESeq2 via PyDESeq2 v0.5.4,^65,66^ with a single factorial model (∼strain + treatment + strain:treatment). Four contrasts were extracted: LPC versus no treatment within each strain and B3 versus WT with and without LPC. For each contrast, genes were ranked by the signed Wald statistic.

GO annotations were compiled from UniProt and supplemented with orthology-based transfer via eggNOG-mapper v2.1.13 (default DIAMOND mode, eggNOG 5.0 database, no taxonomic restriction)^67^ and protein-domain annotation using InterProScan (v5.78-109.0)^68^. GO terms marked obsolete or in the GO Consortium’s “do not annotate” subset was excluded.

Pre-ranked GSEA^69^ was performed using GSEApv v1.3.2 with a weighted statistic (p=1),^70^ 10,000 permutations and gene-sets containing 5-500 represented genes. Gene-set significance was defined as GSEA FDR q value <0.05. The primary analysis used direct, source-assigned GO terms. The analysis was also repeated after propagation through is_a and part_of relationships using otherwise identical parameters. GSEA results for all four comparisons and the ancestor-propagated analysis are provided in Table S3.

### Bioinformatic analyses of FarR variants

A total of 2,042 annotated complete *Staphylococcus aureus* RefSeq assemblies available on NCBI Datasets till Aug. 25^th^, 2026, were retrieved, each with genomic and predicted-protein FASTA files. FarR proteins were initially identified using the NCBI product annotation “fatty acid efflux pump transcriptional regulator FarR.” Candidate sequences were aligned to the 182-amino-acid FarR reference sequence WP_000665238.1 using a Needleman–Wunsch alignment, with match, mismatch, and gap scores of +2, −1, and −2, respectively. Candidates were retained when they were 160–205 amino acids long, had at least 80% reference-normalized identity, and covered at least 90% of the reference sequence. In assemblies without a qualifying annotated FarR protein , other predicted proteins were screened for sequence similarity to the reference.

Remaining unresolved assemblies were searched in all six translated genomic reading frames. Genomically identified sequences were considered complete only when they began with methionine, were at least 180 amino acids long, covered at least 99% of the reference, and met the same identity threshold. This procedure resolved a full-length FarR protein in 2,025 assemblies.

Sequences were projected onto WP_000665238.1 coordinates using an insertion-aware reference alignment. The recurrent glutamate insertion following residue E93 was represented as position 93a, preventing displacement of residue numbering downstream of the insertion. For each reference position, the percentage of assemblies encoding an non-reference amino acid (different from WP_000665238.1) was calculated. The experimentally identified substitutions D94Y, V115I, and E151K were counted only when the exact replacement was present; alternative substitutions at the same position were recorded separately.

### Data Accession

All sequencing reads of this study have been uploaded to the NCBI database with the ProjectID PRJNA1005284 and SampleID SRR25630066-77 (whole genome sequencing results) and SRR39572856-67 (RNA-sequencing results).

## Acknowledgements

The authors acknowledge Luciano Perez Medina Ph.D. from Augusta University for generating the chemical structures of Fig 1A and B. This work was supported by R01AI148366 to K.P. and Z.G. and by the Cecil H. and Ida Green Chair in Systems Biology Science to K.P. Fig 2A was created in BioRender. Wei, Y. (2026) https://BioRender.com/px1r194

## Author contributions

Conceptualization: Y.W. and K.L.P; experiment performance and data analysis: Y.W., Z.G., C.U., S.E.P., J.N., P.M.C., G.C., L.E.T., and G.H.C.; manuscript preparation: Y.W., Z.G., C.U., S.E.P., J.N. and K.L.P.

## Additional Information

The authors declare no known conflicts of interest.

## Reference

1 Xu, J., Lu, D. & Huang, X. Lipid metabolism and metabolites: emerging roles in systemic physiology and metabolic diseases. J Genet Genomics 53, 1143–1158 (2026). 10.1016/j.jgg.2026.01.006

2 Kang, J. H., Toita, R., Kawano, T., Murata, M. & Kano, A. Phospholipids and their metabolites as diagnostic biomarkers of human diseases. Prog Lipid Res 99, 101340 (2025). 10.1016/j.plipres.2025.101340

3 Curran, C. S., Remaley, A. T. & Torabi-Parizi, P. Plasmalogens as biomarkers and therapeutic targets. J Lipid Res 66, 100925 (2025). 10.1016/j.jlr.2025.100925

4 Dorninger, F., Werner, E. R., Berger, J. & Watschinger, K. Regulation of plasmalogen metabolism and traffic in mammals: The fog begins to lift. Front Cell Dev Biol 10, 946393 (2022). 10.3389/fcell.2022.946393

5 Braverman, N. E. & Moser, A. B. Functions of plasmalogen lipids in health and disease. Biochim Biophys Acta 1822, 1442–1452 (2012). 10.1016/j.bbadis.2012.05.008

6 Hughes, B. P. & Frais, F. F. Changes in plasmalogen content of human heart and skeletal muscle with age and development. Nature 215, 993–994 (1967). 10.1038/215993a0

7 Bozelli, J. C., Jr., Azher, S. & Epand, R. M. Plasmalogens and chronic inflammatory diseases. Front Physiol 12, 730829 (2021). 10.3389/fphys.2021.730829

8 Law, S. H. et al. An updated review of lysophosphatidylcholine metabolism in human diseases. Int J Mol Sci 20, 1149–1149 (2019). 10.3390/ijms20051149

9 Zhong, X. et al. Lipidomics based on UHPLC/Q-TOF-MS to characterize lipid metabolic profiling in patients with newly diagnosed type 2 diabetes mellitus with dyslipidemia. Heliyon 10, e26326 (2024). 10.1016/j.heliyon.2024.e26326

10 Yea, K. et al. Lysophosphatidylcholine activates adipocyte glucose uptake and lowers blood glucose levels in murine models of diabetes. J Biol Chem 284, 33833–33840 (2009). 10.1074/jbc.M109.024869

11 Barber, M. N. et al. Plasma lysophosphatidylcholine levels are reduced in obesity and type 2 diabetes. PLoS One 7, e41456 (2012). 10.1371/journal.pone.0041456

12 Chávez-Reyes, J. et al. Susceptibility for some infectious diseases in patients with diabetes: the key role of glycemia. Front Public Health 9, 559595 (2021). 10.3389/fpubh.2021.559595

13 Kim, T. & Choi, S. H. Diabetes mellitus and infectious diseases: current evidence and clinical implications. Diabetes Metab J 49, 915–933 (2025). 10.4093/dmj.2025.0508

14 Miyazaki, H. et al. Antimicrobial effects of lysophosphatidylcholine on methicillin-resistant *Staphylococcus aureus*. Ther Adv Infect Dis 4, 89–94 (2017). 10.1177/2049936117714920

15 Riessberger-Gallé, U., Hernández-López, J., Rechberger, G., Crailsheim, K. & Schuehly, W. Lysophosphatidylcholine acts in the constitutive immune defence against American foulbrood in adult honeybees. Sci Rep 6, 30699 (2016). 10.1038/srep30699

16 Parra-Millán, R. et al. Impact of the immune response modification by lysophosphatidylcholine in the efficacy of antibiotic therapy of experimental models of peritoneal sepsis and pneumonia by *Pseudomonas aeruginosa*: LPC therapeutic effect in combined therapy. Enfermedades Infecciosas y Microbiologia Clinica (2020). 10.1016/j.eimc.2020.06.002

17 Tsuchido, T. Induction of cell autolysis of *Bacillus subtilis* with lysophosphatidylcholine. Applied Microbiology and Biotechnology 41, 106–109 (1994). 10.1007/BF00166090

18 Yan, J. J. et al. Therapeutic effects of lysophosphatidylcholine in experimental sepsis. Nature Medicine 10, 161–167 (2004). 10.1038/nm989

19 Parra Millán, R., et al. Efficacy of lysophosphatidylcholine in combination with antimicrobial agents against *Acinetobacter baumannii* in experimental murine peritoneal sepsis and pneumonia models. Antimicrob Agents Chemother 60, 4464–4470 (2016). 10.1128/AAC.02708-15

20 Kondo, E. & Kanai, K. Mechanism of bactericidal activity of lysolecithin and its biological implication. Jpn J Med Sci Biol 38, 181–194 (1985). 10.7883/yoken1952.38.181

21 Yoon, B. K., Jackman, J. A., Valle-Gonzalez, E. R. & Cho, N. J. Antibacterial free fatty acids and monoglycerides: biological activities, experimental testing, and therapeutic applications. Int J Mol Sci 19 (2018). 10.3390/ijms19041114

22 Parsons, J. B., Yao, J., Frank, M. W., Jackson, P. & Rock, C. O. Membrane disruption by antimicrobial fatty acids releases low-molecular-weight proteins from *Staphylococcus aureus*. J Bacteriol 194, 5294–5304 (2012). 10.1128/JB.00743-12

23 Carson, D. D. & Daneo-Moore, L. Effects of fatty acids on lysis of *Streptococcus faecalis*. J Bacteriol 141, 1122–1126 (1980). 10.1128/jb.141.3.1122-1126.1980

24 Galbraith, H., Miller, T. B., Paton, A. M. & Thompson, J. K. Antibacterial activity of long chain fatty acids and the reversal with calcium, magnesium, ergocalciferol and cholesterol. J Appl Bacteriol 34, 803–813 (1971). 10.1111/j.1365-2672.1971.tb01019.x

25 Kabara, J. J., Swieczkowski, D. M., Conley, A. J. & Truant, J. P. Fatty acids and derivatives as antimicrobial agents. Antimicrob Agents Chemother 2, 23–28 (1972). 10.1128/AAC.2.1.23

26 Kengmo Tchoupa, A., Eijkelkamp, B. A. & Peschel, A. Bacterial adaptation strategies to host-derived fatty acids. Trends in Microbiology 30, 241–253 (2022). 10.1016/j.tim.2021.06.002

27 Parsons, J. B. et al. Identification of a two-component fatty acid kinase responsible for host fatty acid incorporation by *Staphylococcus aureus*. Proc Natl Acad Sci U S A 111, 10532–10537 (2014). 10.1073/pnas.1408797111

28 Gullett, J. M., Cuypers, M. G., Frank, M. W., White, S. W. & Rock, C. O. A fatty acid-binding protein of *Streptococcus pneumoniae* facilitates the acquisition of host polyunsaturated fatty acids. J Biol Chem 294, 16416–16428 (2019). 10.1074/jbc.RA119.010659

29 Lambert, C. et al. A *Streptococcus pyogenes* DegV protein regulates the membrane lipid content and limits the formation of extracellular vesicles. PLoS One 18, e0284402 (2023). 10.1371/journal.pone.0284402

30 Zou, Q., Dong, H. & Cronan, J. E. The enteric bacterium *Enterococcus faecalis* elongates and incorporates exogenous short and medium chain fatty acids into membrane lipids. Mol Microbiol 122, 757–771 (2024). 10.1111/mmi.15322

31 Alnaseri, H. et al. Inducible expression of a resistance-nodulation-division-type efflux pump in *Staphylococcus aureus* provides resistance to linoleic and arachidonic acids. J Bacteriol 197, 1893–1905 (2015). 10.1128/JB.02607-14

32 Schmiel, D. H. & Miller, V. L. Bacterial phospholipases and pathogenesis. Microbes Infect 1, 1103–1112 (1999). 10.1016/s1286-4579(99)00205-1

33 Sakasegawa, S. I., Maeba, R., Murayama, K., Matsumoto, H. & Sugimori, D. Hydrolysis of plasmalogen by phospholipase A1 from Streptomyces albidoflavus for early detection of dementia and arteriosclerosis. Biotechnol Lett 38, 109–116 (2016). 10.1007/s10529-015-1955-5

34 Hayashi, D., Mouchlis, V. D. & Dennis, E. A. Each phospholipase A(2) type exhibits distinct selectivity toward sn-1 ester, alkyl ether, and vinyl ether phospholipids. Biochim Biophys Acta Mol Cell Biol Lipids 1867, 159067 (2022). 10.1016/j.bbalip.2021.159067

35 Wu, L. C. et al. Purification, identification, and cloning of lysoplasmalogenase, the enzyme that catalyzes hydrolysis of the vinyl ether bond of lysoplasmalogen. J Biol Chem 286, 24916–24930 (2011). 10.1074/jbc.M111.247163

36 van Hensbergen, V. P., Wu, Y., van Sorge, N. M. & Touqui, L. Type IIA Secreted Phospholipase A2 in Host Defense against Bacterial Infections. Trends Immunol 41, 313–326 (2020). 10.1016/j.it.2020.02.003

37 Joyce, L. R., Guan, Z. & Palmer, K. L. Phosphatidylcholine Biosynthesis in Mitis Group Streptococci via Host Metabolite Scavenging. J Bacteriol 201 (2019). 10.1128/JB.00495-19

38 Joyce, L. R., Guan, Z. & Palmer, K. L. *Streptococcus pneumoniae*, *S. pyogenes* and *S. agalactiae* membrane phospholipid remodelling in response to human serum. Microbiology (Reading) 167 (2021). 10.1099/mic.0.001048

39 Moser, R., Aktas, M. & Narberhaus, F. Phosphatidylcholine biosynthesis in Xanthomonas campestris via a yeast-like acylation pathway. Mol Microbiol 91, 736–750 (2014). 10.1111/mmi.12492

40 Kleetz, J., Vasilopoulos, G., Czolkoss, S., Aktas, M. & Narberhaus, F. Recombinant and endogenous ways to produce methylated phospholipids in *Escherichia coli*. Appl Microbiol Biotechnol 105, 8837–8851 (2021). 10.1007/s00253-021-11654-8

41 Jackson, D. R. et al. Plasmalogen Biosynthesis by Anaerobic Bacteria: Identification of a Two-Gene Operon Responsible for Plasmalogen Production in *Clostridium perfringens*. ACS Chem Biol 16, 6–13 (2021). 10.1021/acschembio.0c00673

42 Wishart, D. S. et al. HMDB 5.0: the Human Metabolome Database for 2022. Nucleic Acids Res 50, D622–D631 (2022). 10.1093/nar/gkab1062

43 Giallourou, N. et al. Characterizing the breast cancer lipidome and its interaction with the tissue microbiota. Commun Biol 4, 1229 (2021). 10.1038/s42003-021-02710-0

44 Iwasaki, T. et al. Plasma Lysophosphatidylcholine Levels Correlate with Prognosis and Immunotherapy Response in Squamous Cell Carcinoma. Int J Mol Sci 26 (2025). 10.3390/ijms26157528

45 Kalia, V. et al. Lysophosphatidylcholines are associated with amyloidosis in early stages of Alzheimer’s disease. Nat Aging 6, 221–234 (2026). 10.1038/s43587-025-01025-7

46 Nguyen, M. T. et al. Inactivation of *farR* causes high rhodomyrtone resistance and increased pathogenicity in *Staphylococcus aureus*. Front Microbiol 10, 1–14 (2019). 10.3389/fmicb.2019.01157

47 Yao, J. & Rock, C. O. Bacterial fatty acid metabolism in modern antibiotic discovery. Biochim Biophys Acta Mol Cell Biol Lipids 1862, 1300–1309 (2017). 10.1016/j.bbalip.2016.09.014

48 Howden, B. P. et al. Evolution of multidrug resistance during *Staphylococcus aureus* infection involves mutation of the essential two component regulator WalKR. PLoS Pathogens 7, e1002359 (2011). 10.1371/journal.ppat.1002359

49 Chen, Y., Moran, J. C., Campbell-Lee, S. & Horsburgh, M. J. Transcriptomic Responses and Survival Mechanisms of Staphylococci to the Antimicrobial Skin Lipid Sphingosine. Antimicrob Agents Chemother 66, e0056921 (2022). 10.1128/aac.00569-21

50 Huang, L. et al. Molecular Basis of Rhodomyrtone Resistance in *Staphylococcus aureus*. mBio 13, e0383321 (2022). 10.1128/MBIO.03833-21

51 Lin, Y. T. et al. Emergence of a small colony variant of vancomycin-intermediate *Staphylococcus aureus* in a patient with septic arthritis during long-term treatment with daptomycin. J Antimicrob Chemother 71, 1807–1814 (2016). 10.1093/jac/dkw060

52 Bonn, C. M. et al. Repeated emergence of variant TetR family regulator, FarR, and increased resistance to antimicrobial unsaturated fatty acid among Clonal Complex 5 methicillin-resistant *Staphylococcus aureus*. Antimicrob Agents Chemother 67, e0074922 (2023). 10.1128/aac.00749-22

53. *European Committee on Antimicrobial Susceptibility Testing. Breakpoint tables for interpretation of MICs and zone diameters. Version 16.1*, <https://www.eucast.org> (2026).

54 Zhang, L., Gray, L., Novick, R. P. & Ji, G. Transmembrane topology of AgrB, the protein involved in the post-translational modification of AgrD in *Staphylococcus aureus*. J Biol Chem 277, 34736–34742 (2002). 10.1074/jbc.M205367200

55 Kinkel, T. L., Roux, C. M., Dunman, P. M. & Fang, F. C. The *Staphylococcus aureus* SrrAB two-component system promotes resistance to nitrosative stress and hypoxia. mBio 4, e00696–00613 (2013). 10.1128/mBio.00696-13

56 Majerczyk, C. D. et al. Direct targets of CodY in *Staphylococcus aureus*. J Bacteriol 192, 2861–2877 (2010). 10.1128/JB.00220-10

57 Gründling, A. & Schneewind, O. Synthesis of glycerol phosphate lipoteichoic acid in *Staphylococcus aureus*. Proceedings of the National Academy of Sciences 104, 8478–8483 (2007). 10.1073/pnas.0701821104

58 Tran, T. T., Miller, W. R., Shamoo, Y. & Arias, C. A. Targeting cell membrane adaptation as a novel antimicrobial strategy. Curr Opin Microbiol 33, 91–96 (2016). 10.1016/j.mib.2016.07.002

59 Shi, Y. et al. Structure and mechanism for streptococcal fatty acid kinase (Fak) system dedicated to host fatty acid scavenging. Sci Adv 8, eabq3944 (2022). 10.1126/sciadv.abq3944

60 Chacko, K. I. et al. Genetic basis of emerging vancomycin, linezolid, and daptomycin heteroresistance in a case of persistent *Enterococcus faecium* bacteremia. Antimicrob Agents Chemother 62 (2018). 10.1128/AAC.02007-17

61 Van De Rijn, I. Quantitative analysis of cell walls of nutritionally variant streptococci grown under various growth conditions. Infection and Immunity 49, 518–522 (1985). 10.1128/iai.49.3.518-522.1985

62 Wei, Y., Joyce, L. R., Wall, A. M., Guan, Z. & Palmer, K. L. *Streptococcus pneumoniae*, *S. mitis*, and *S. oralis* produce a phosphatidylglycerol-dependent, *ltaS*-independent glycerophosphate-linked glycolipid. mSphere 6, e01099–01020 (2021). 10.1128/mSphere.01099-20

63 Wei, Y., Sturges, C. I. & Palmer, K. L. Human serum supplementation promotes *Streptococcus mitis* growth and induces specific transcriptomic responses. Microbiology Spectrum **Apr**, e0512922–e0512922 (2023). 10.1128/spectrum.05129-22

64 Untergasser, A. et al. Primer3--new capabilities and interfaces. Nucleic Acids Res 40, e115 (2012). 10.1093/nar/gks596

65 Love, M. I., Huber, W. & Anders, S. Moderated estimation of fold change and dispersion for RNA-seq data with DESeq2. Genome Biol 15, 550 (2014). 10.1186/s13059-014-0550-8

66 Muzellec, B., Telenczuk, M., Cabeli, V. & Andreux, M. PyDESeq2: a python package for bulk RNA-seq differential expression analysis. Bioinformatics 39 (2023). 10.1093/bioinformatics/btad547

67 Cantalapiedra, C. P., Hernandez-Plaza, A., Letunic, I., Bork, P. & Huerta-Cepas, J. eggNOG-mapper v2: Functional Annotation, Orthology Assignments, and Domain Prediction at the Metagenomic Scale. Mol Biol Evol 38, 5825–5829 (2021). 10.1093/molbev/msab293

68 Jones, P. et al. InterProScan 5: genome-scale protein function classification. Bioinformatics 30, 1236–1240 (2014). 10.1093/bioinformatics/btu031

69 Subramanian, A. et al. Gene set enrichment analysis: a knowledge-based approach for interpreting genome-wide expression profiles. Proc Natl Acad Sci U S A 102, 15545–15550 (2005). 10.1073/pnas.0506580102

70 Fang, Z., Liu, X. & Peltz, G. GSEApy: a comprehensive package for performing gene set enrichment analysis in Python. Bioinformatics 39 (2023). 10.1093/bioinformatics/btac757

