## Supplemental materials for "Gram-positive bacteria have different susceptibilities to the host-derived antimicrobial lysophosphatidylcholine and adapt to lysophosphatidylcholine-induced stress via distinct mechanisms"

**Supplementary materials**:

**Materials and methods**:

*Lipidomic Analysis by Normal Phase LC/MS/MS*

Normal phase LC was performed on an Agilent 1200 Quaternary LC system equipped with an Ascentis Silica HPLC column, 5 μm, 25 cm x 2.1 mm (Sigma-Aldrich, St. Louis, MO). Mobile phase A consisted of chloroform/methanol/aqueous ammonium hydroxide (800:195:5, v/v); mobile phase B consisted of chloroform/methanol/water/aqueous ammonium hydroxide (600:340:50:5, v/v); mobile phase C consisted of chloroform/methanol/water/aqueous ammonium hydroxide (450:450:95:5, v/v). The elution program consisted of the following: 100% mobile phase A was held isocratically for 2 min and then linearly increased to 100% mobile phase B over 8 min and held at 100% B for 5 min. The LC gradient was then changed to 100% mobile phase C over 1 min and held at 100% C for 3 min, and finally returned to 100% A over 0.5 min and held at 100% A for 3 min. The LC eluent (with a total flow rate of 300 μL/min) was introduced into the ESI source of a high resolution TripleTOF5600 mass spectrometer (Sciex, Framingham, MA). Instrumental settings for negative ion ESI and MS/MS analysis of lipid species were as follows: IS= -4500 V; CUR= 20 psi; GSI= 20 psi; DP= -55 V; and FP= -150 V. The MS/MS analysis used nitrogen as the collision gas. Data analysis was performed using Analyst TF1.5 software (Sciex, Framingham, MA).

**Result Figures and Tables:**

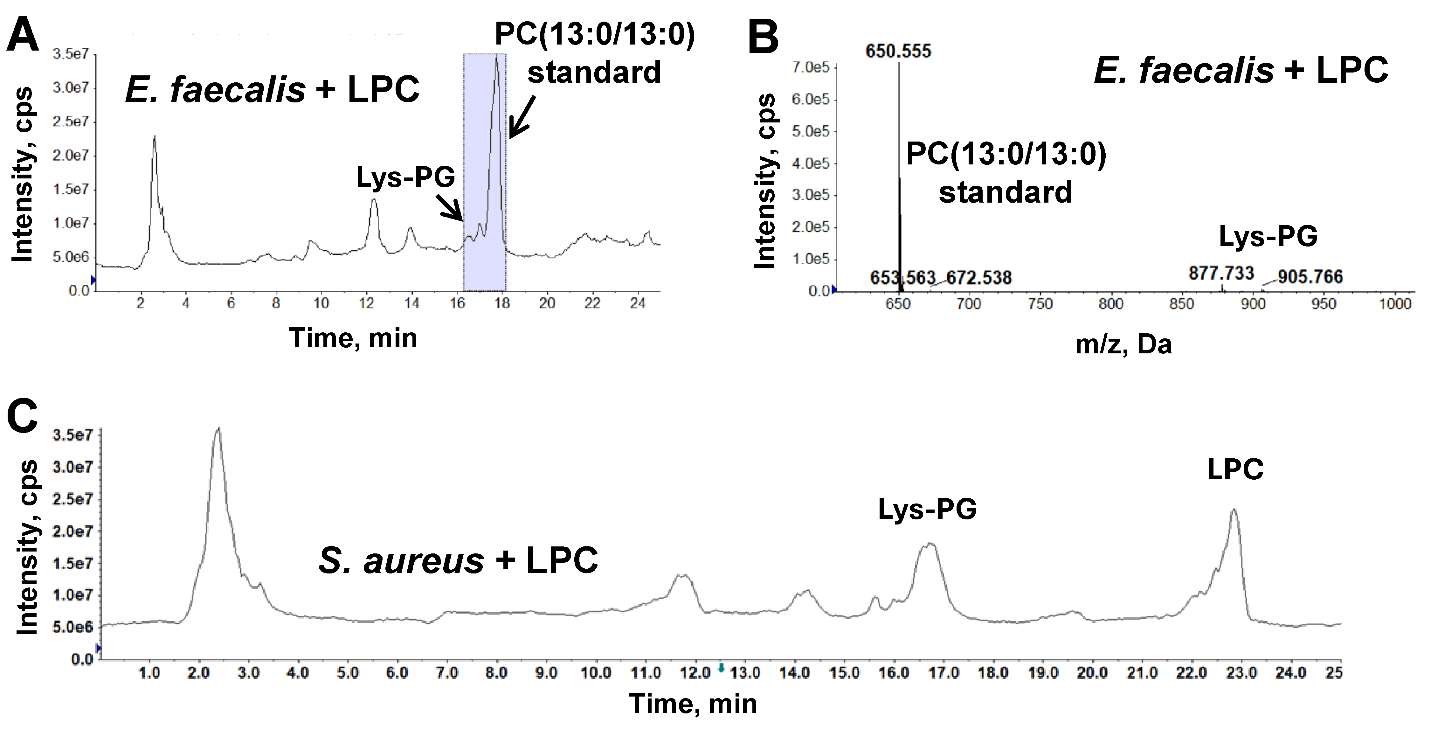

**Fig S1**: **Neither *S. aureus* nor *E. faecalis* scavenges LPC for the synthesis of PC.** A. Total ion chromatogram of LC/MS of lipid extract of *E. faecalis* grown to stationary phase in CDM supplemented with 50 μM LPC. B. Mass spectrum (averaged from retention time 16-18 minutes) showing PC (13:0/13:0) spike-in standard and the bacterial-synthesized lipid Lys-PG. C. Total ion chromatogram of LC/MS of lipid extract of *S. aureus* grown to stationary phase in chemically defined media (CDM) supplemented with 50 μM LPC. Abbreviations: PC, phosphatidylcholine; PG, phosphatidylglycerol; Lys-PG, lysyl-phosphatidylglycerol; LPC, lysophosphatidylcholine.

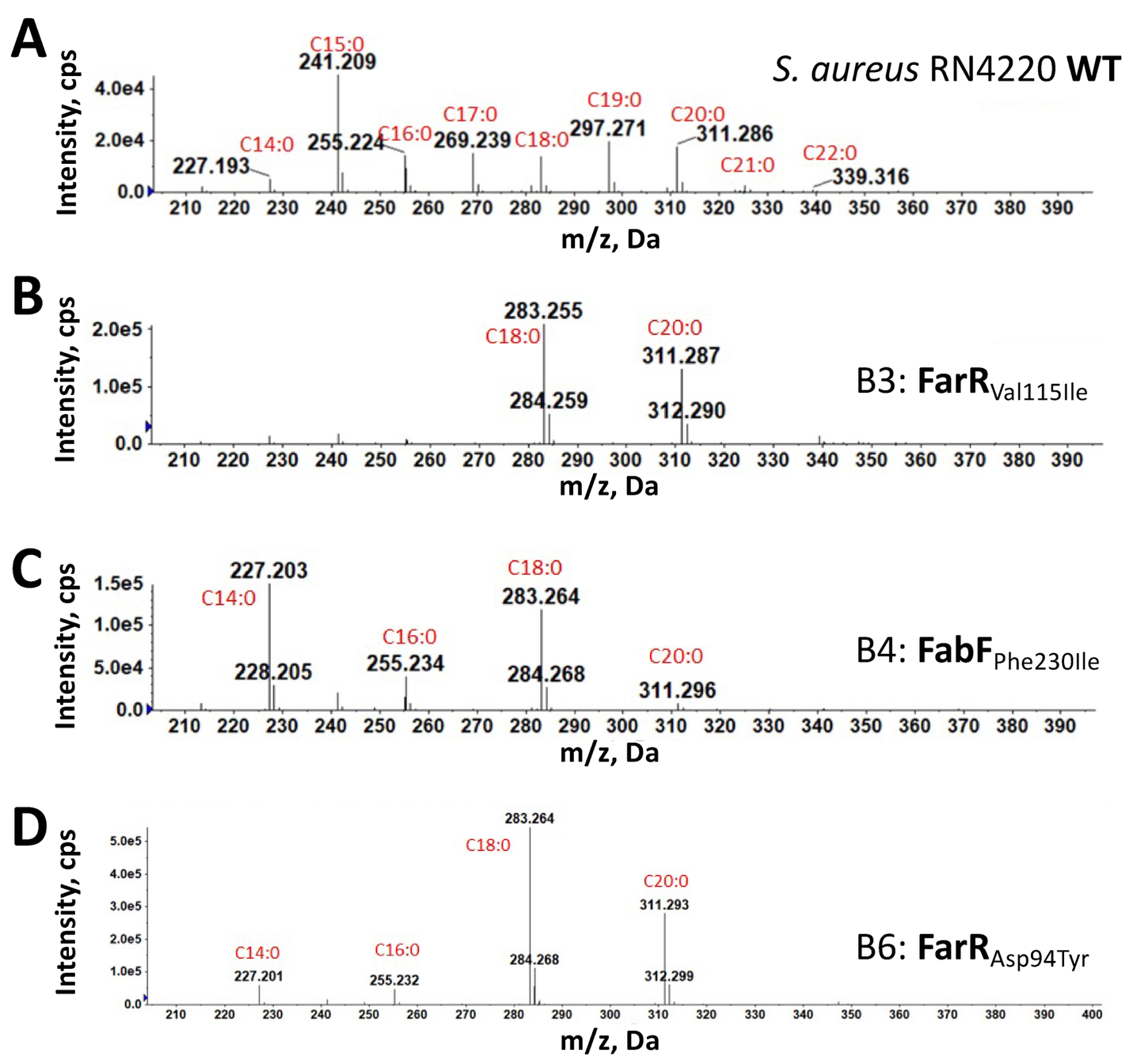

**Fig S2**: Negative ion mass spectra of free fatty acids of *S. aureus* RN4220 wildtype (WT) and mutants.

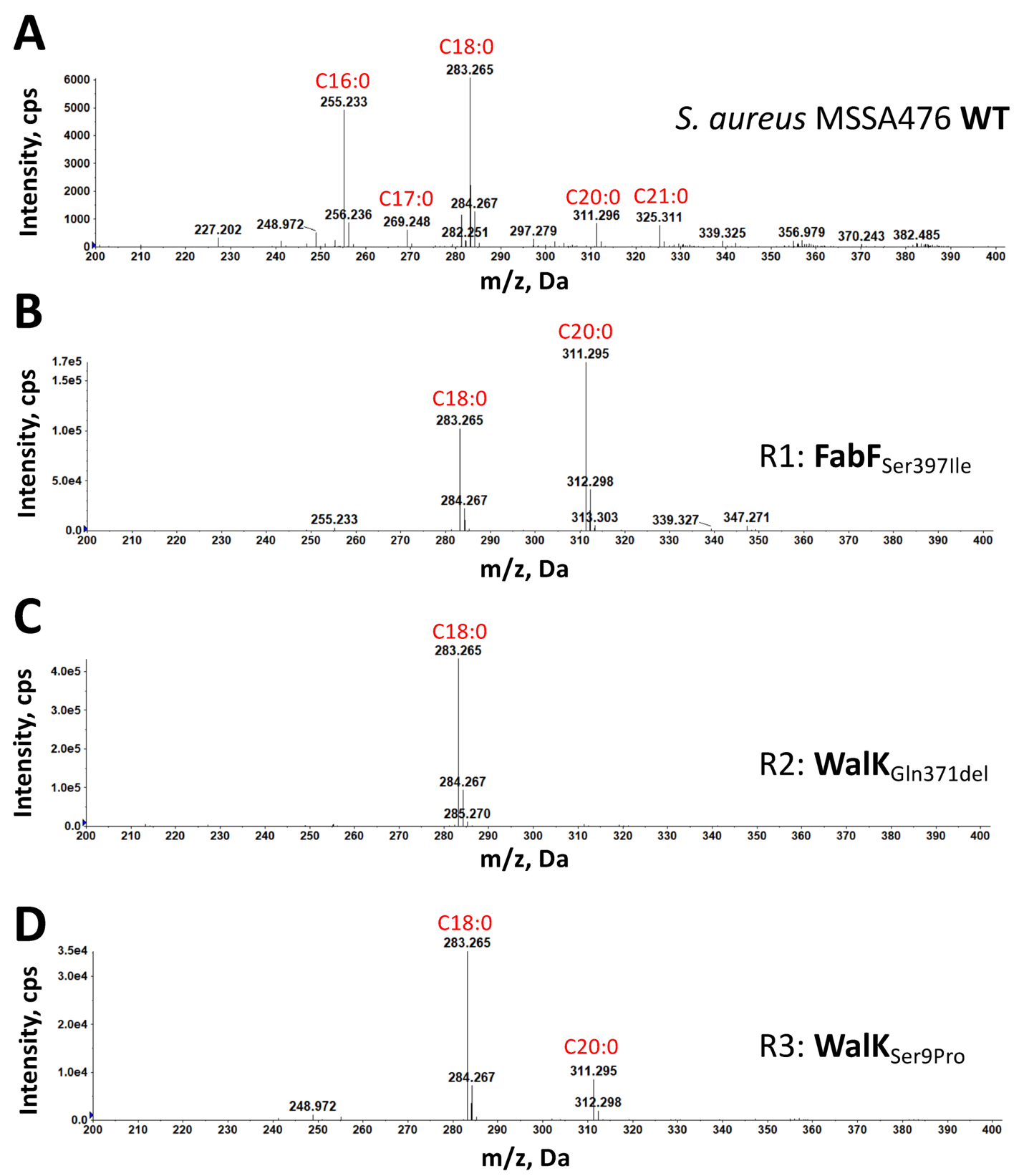

**Fig S3**: Negative ion mass spectra of free fatty acids of *S. aureus* MSSA476 wildtype (WT) and mutants.

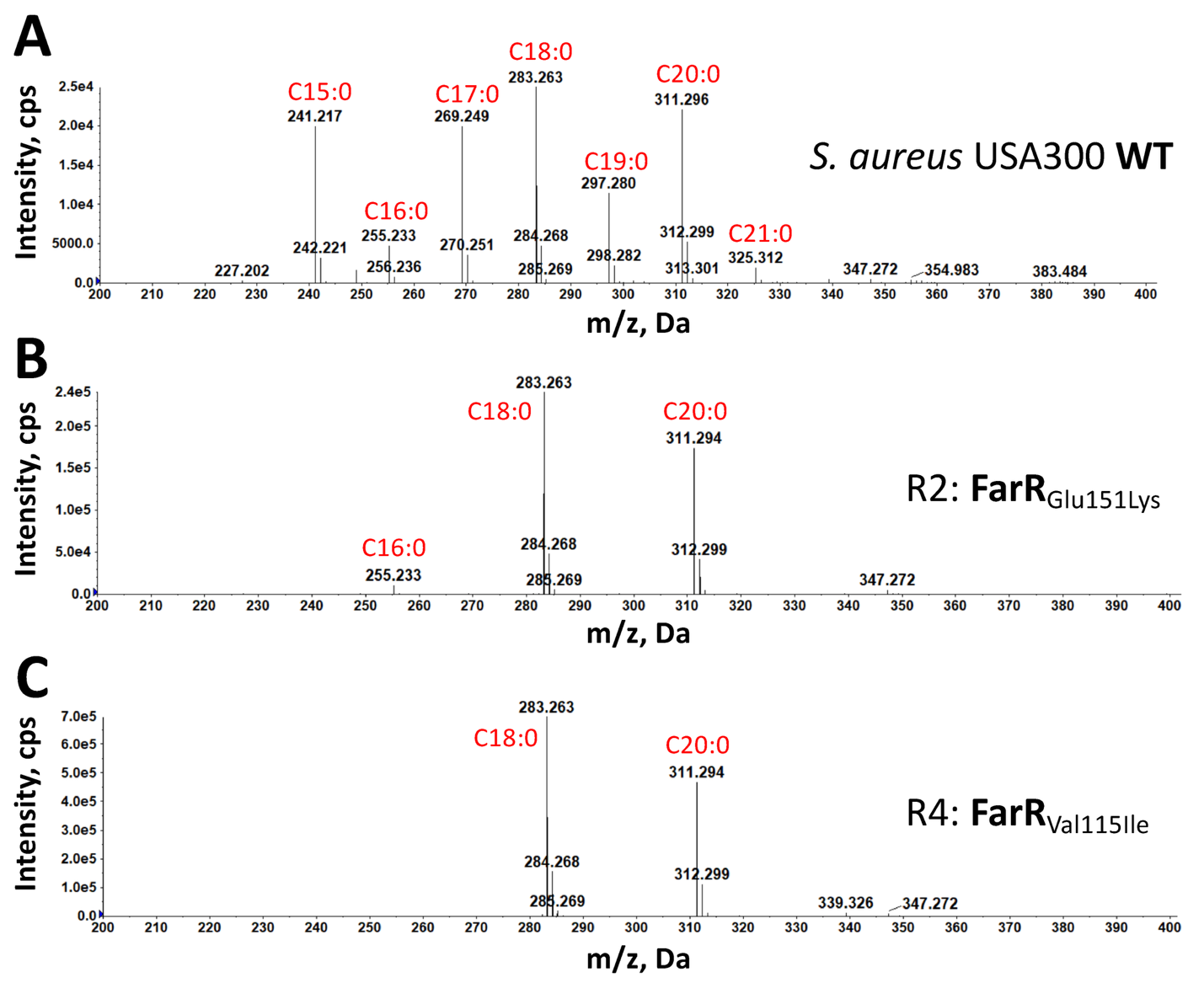

**Fig S4**: Negative ion mass spectra of free fatty acids of *S. aureus* USA300 wildtype (WT) and mutants.

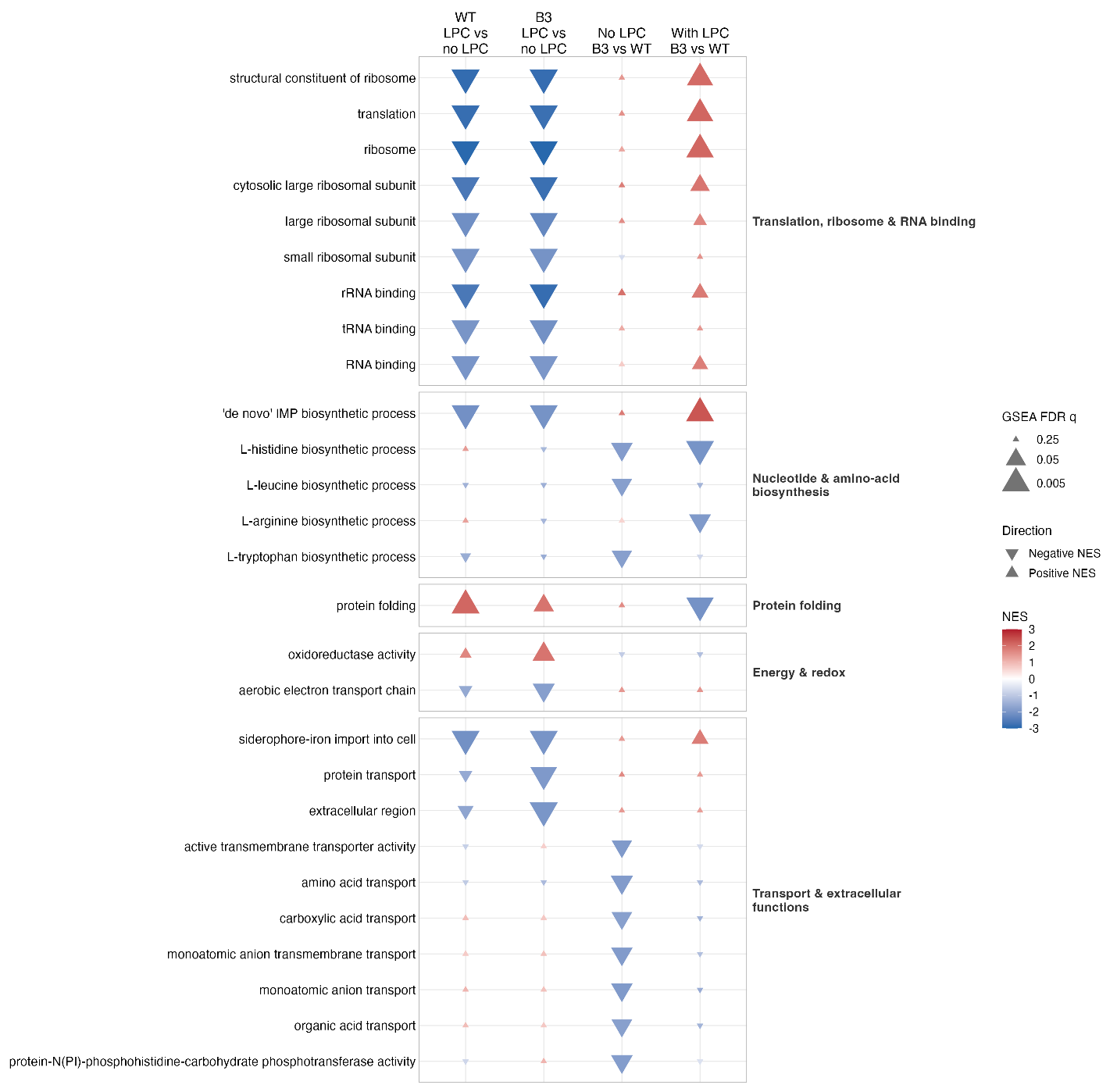

**Fig S5: GO enrichment across strain and LPC comparisons.** Direct GO terms significant in at least one comparison (GSEA FDR q<0.05) are shown. Upward and downward triangles indicate positive and negative NES, respectively; positive NES indicates enrichment with LPC in treatment comparisons and enrichment in B3 in strain comparisons. Color represents NES, and triangle size represents the GSEA FDR q value, with larger triangles indicating smaller q values.

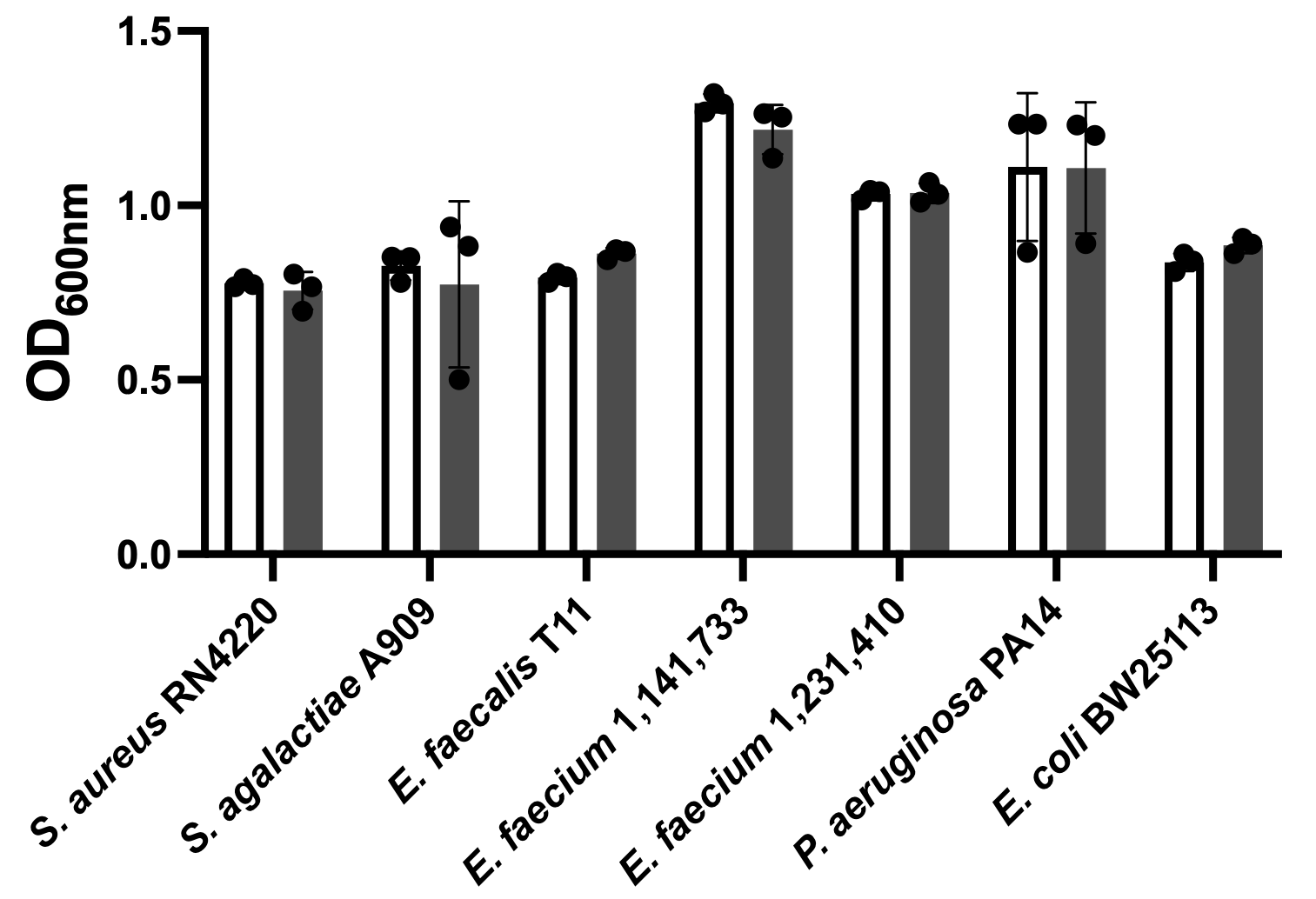

**Fig S6**: **Methanol has no adverse effects on the growth of selected bacteria**. Bacterial cultures (inoculated from a single colony) were diluted to an OD_600nm_ of 0.1 in chemically defined medium either supplemented with methanol to 0.2% (v/v) (solid grey bar), or without the supplementation (empty bar). Bacterial growth was measured after 24 hours incubation at 37°C. Each data point represents a biologically independent replicate. Statistical analyses were performed with student *t*-test to compare the growth of a bacterium in different cultivation media. None of the tested bacterial strains show significant (P-value < 0.05) growth differences between the conditions.

**Table S1: R^2^ values of the linear regression analyses of RNA-sequencing RPKMs**

| R^2^ values | B3 in CDM | B3 in CDM with LPC | WT in CDM | WT in CDM with LPC |
| --- | --- | --- | --- | --- |
| Replicate 1 & 2 | 0.99277 | 0.99244 | 0.95286 | 0.98919 |
| Replicate 1 & 3 | 0.97402 | 0.99359 | 0.97959 | 0.98264 |
| Replicate 2 & 3 | 0.96877 | 0.97756 | 0.96656 | 0.98906 |

**Table S2: RNA sequencing results** (Excel file).

**Table S3: Gene set enrichment analytical results** (Excel file).
